# High temporal resolution systems profiling reveals distinct patterns of interferon response after Covid-19 mRNA vaccination and SARS-CoV2 infection

**DOI:** 10.1101/2021.12.12.472257

**Authors:** Darawan Rinchai, Sara Deola, Gabriele Zoppoli, Basirudeen Syed Ahamed Kabeer, Sara Taleb, Igor Pavlovski, Selma Maacha, Giusy Gentilcore, Mohammed Toufiq, Lisa Mathew, Li Liu, Fazulur Rehaman Vempalli, Ghada Mubarak, Stephan Lorenz, Irene Sivieri, Gabriella Cirmena, Chiara Dentone, Paola Cuccarolo, Daniele Roberto Giacobbe, Federico Baldi, Alberto Garbarino, Benedetta Cigolini, Paolo Cremonesi, Michele Bedognetti, Alberto Ballestrero, Matteo Bassetti, Boris P. Hejblum, Tracy Augustine, Nicholas Van Panhuys, Rodolphe Thiebaut, Ricardo Branco, Tracey Chew, Maryam Shojaei, Kirsty Short, Carl Feng, PREDICT-19 consortium, Susu M. Zughaier, Andrea De Maria, Benjamin Tang, Ali Ait Hssain, Davide Bedognetti, Jean-Charles Grivel, Damien Chaussabel

## Abstract

Knowledge of the mechanisms underpinning the development of protective immunity conferred by mRNA vaccines is fragmentary. Here we investigated responses to COVID-19 mRNA vaccination via ultra-low-volume sampling and high-temporal-resolution transcriptome profiling (23 subjects across 22 timepoints, and with 117 COVID-19 patients used as comparators). There were marked differences in the timing and amplitude of the responses to the priming and booster doses. Notably, we identified two distinct interferon signatures. The first signature (A28/S1) was robustly induced both post-prime and post-boost and in both cases correlated with the subsequent development of antibody responses. In contrast, the second interferon signature (A28/S2) was robustly induced only post-boost, where it coincided with a transient inflammation peak. In COVID19 patients, a distinct phenotype dominated by A28/S2 was associated with longer duration of intensive care. In summary, high-temporal-resolution transcriptomic permitted the identification of post-vaccination phenotypes that are determinants of the course of COVID-19 disease.

## INTRODUCTION

COVID-19 vaccines are critical to the ongoing efforts to control the SARS-CoV-2 coronavirus pandemic. To date, nine vaccines have received some form of approval for use in humans, and phase III trials are ongoing for an additional 11 vaccines (1). Notable differences exist among the vaccine products in terms of their design and the levels of protection they confer, as well as the type, incidence, and severity of adverse events they may elicit. Gaining a comprehensive understanding of the immunological factors underpinning the different responses to various vaccines is a major endeavor. Yet, this knowledge is necessary for guiding timely decisions to modulate vaccination protocols (e.g., the use of different types of vaccines for the priming and booster doses). This information may also assist in matching of individuals with the growing number of available vaccines based on their demographics, health status, or any other relevant clinical/molecular phenotypes.

Blood transcriptome profiling measures the abundance of transcripts in whole blood and on a system-wide scale. It was previously employed to comprehensively profile the immune responses elicited by vaccines (2, 3). Notably, this approach identified innate immune signatures arising within hours after administering vaccines (4). In a recently published report, Arunachalam et al. described the blood transcriptome profiles measured following the administration of the BNT162b2 mRNA COVID-19 vaccine (5). They reported the presence of an interferon (IFN) signature one day after the priming vaccination that was no longer detectable on day 7. They further found a more comprehensive IFN/inflammatory signature to be present 1 day after administering the booster dose. However, the sampling schedule employed in this study was relatively sparse. And the sample collection time points commonly selected in systems vaccinology studies are based on kinetics established for more conventional vaccines – with sampling at days 1 and 7 often selected since they correspond to the peaks of the innate and adaptive immune responses elicited for instance by the influenza or pneumococcal vaccines (6). However, the precise kinetics of the immune response elicited by mRNA vaccines remains to be established. In the present study we endeavored to profile the blood transcriptome of individuals prior to the administration of the first dose of COVID-19 mRNA vaccine and for the following 9 consecutive days. Subjects also collecting samples for deep serological profiling at three time points. The same sampling and profiling schedule was repeated to assess the response to the second dose of the vaccine. To achieve this, we have adopted a ultra-low volume sampling procedure consisting in the self-collection of few drops of blood (50 ul) by fingerstick (7).

Together, this work permitted the precise delineation of a well-orchestrated immune response to COVID-19 mRNA vaccines and identified marked differences in the magnitude, nature, and timing of the transcriptional signatures elicited by prime and boost vaccination. Most notably, differences in temporal patterns of responsiveness revealed distinct components of the interferon response, which is known to play a key role in controlling SARS-CoV-2 infection (8) and was also found here to associate with the subsequent development of the antibody response post-vaccination.

## RESULTS

### Study design, implementation, and serological profiling

We successfully recruited a cohort of volunteers and implemented a high-frequency sampling protocol. This permitted to ascertain the response to the first and second dose of COVID-19 vaccines at 10 consecutive daily timepoints: immediately before vaccination and for 9 days after. We collected samples for serological profiling at three time points: before vaccination and on days 7 and 14 post-vaccination (**Figure 1A**). We implemented a self-sampling blood collection protocol so that subjects could extract small volumes (50 µl) of RNA-stabilized blood at the required frequency (the approach is described in the Methods section and an earlier publication (7)). RNA sequencing profiles were generated using a cost-effective 3ʹ- biased library preparation protocol (Lexogen QuantSeq), which is optimized for low amount of RNA input. We generated COVID-19-specific antibody profiles from capillary blood samples collected by Volumetric Absorptive Micro Sampling analyzed using a multiplexed Bead array established by our team (see Methods for details). Overall, 23 subjects were enrolled in the study, and the characteristics of this cohort are reported in **Table 1**. They received either two doses of the Pfizer/BioNTech mRNA vaccine (BNT162b2, N = 19) or two doses of the Moderna mRNA vaccine (N = 4). Among those 23 subjects, six had recovered from COVID-19 in the months preceding the administration of the first vaccine dose. In total 440 RNA sequencing profiles were generated, and this extensive dataset was shared publicly in GEO with the accession number GSE190001. The serological profiles included reactivity to a stabilized trimer of Spike protein, the spike protein, its receptor-binding domain, the Nucleo and Envelope proteins, of SARS-CoV2, and the subunit S1 of SARS spike protein. The data are provided in **Supplementary File 1**. The seroreactivity to each of these antigens was dissected by measuring the total IgG, total IgA, and IgM, as well as the finer-scale IgG and IgA subtypes. Serological profiling data showed a rise in the levels of antibodies in the plasma of the subjects post-vaccination (**Figure 1B**), and this included antibodies specific for the SARS-CoV-2 Spike protein, which is targeted by COVID-19 vaccines. No responses to the Envelope protein were detected. Some cross-reactivity was observed with the SARS Spike protein. Notably, higher antibody levels were induced after the first dose in individuals who had been previously infected with the virus (**Figure 1B-C**).

**Figure 1:**
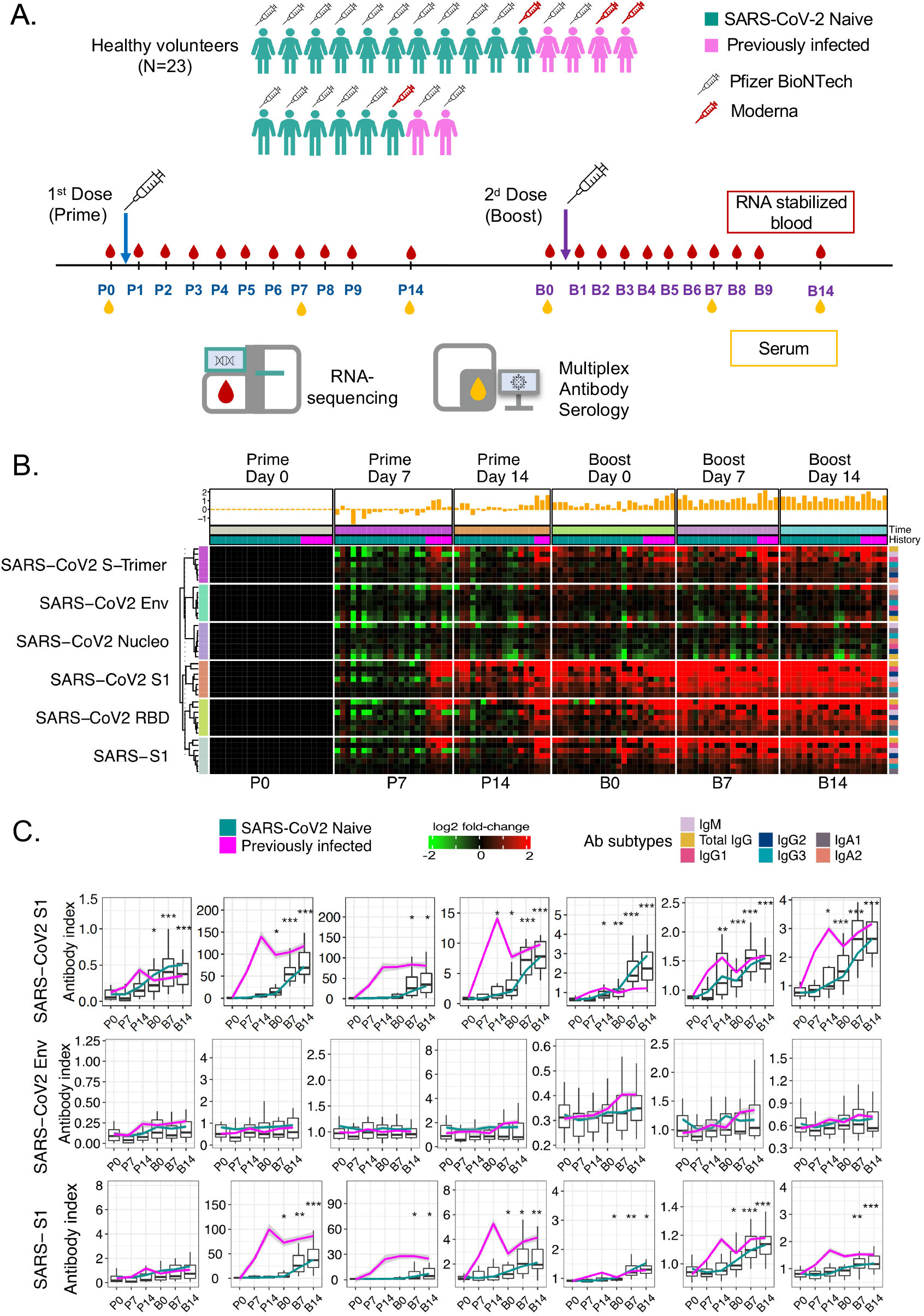
Antibody response to COVID-19 mRNA vaccination. (A) Schematic representation of the study design. (B) The heatmap represents changes in abundance of antibodies specific to several SARS-CoV-2 antigens and control antigens relative to pre-vaccination levels. Red indicates a relative increase, and green indicates a relative decrease in abundance. Columns represent subjects arranged by timepoint and have colored tracks at the top indicating whether the subjects were previously infected with SARS-CoV-2 or not. The histogram above represents the average log2 fold-change over baseline for a given column. The rows represent antibody reactivities arranged by antigen specificity. The different rows represent the isotypes of reactive antibodies, arranged according to the color legend specified below the heat map. (C) Changes in antibody levels expressed as an “antibody index” are shown on the box plots, each corresponding to a given antibody type of a given specificity. Lines indicate changes for individuals previously infected with SARS-CoV-2 and who had recovered (in pink) and for naïve individuals (in green). Centerlines, box limits, and whiskers represent the median, interquartile range, and 1.5x interquartile range, respectively. Multiple pairwise tests (paired t-test) were performed comparing antibodies levels to baseline (D0). Asterisks: * represent p < .01, **represent p < .001, *** represent p < .0001.

**Table 1:** Subject characteristics.

| Patient ID | Vaccine name | Gender | Age | Ethnicity | Previous COVID-19 | Underlying disease | Drugs | Symptoms at prime (Type) | Symptoms at prime (Grade) | Symptoms at boost (Type) | Symptoms at boost (Grade) |
| --- | --- | --- | --- | --- | --- | --- | --- | --- | --- | --- | --- |
| PZB1 | Pfizer Biontech | Female | 38 | Asian | Yes | No | no | Myalgia | G1 | Fever/Myalgia | G1 |
| PZB2 | Pfizer Biontech | Male | 47 | Caucasian | Yes | No | no | Myalgia | G1 | Myalgia | G1 |
| PZB3 | Pfizer Biontech | Male | 57 | Caucasian | Yes | T2D | Metformin, Insulin | Myalgia | G1 | Myalgia | G1 |
| PZB4 | Pfizer Biontech | Female | 34 | Indian | No | No | no | None | NA | Chills/Insomnia/Headache/Myalgia/Fatigue | G3 |
| PZB5 | Pfizer Biontech | Male | 38 | Indian | No | No | no | Myalgia | G1 | Myalgia | G1 |
| PZB6 | Pfizer Biontech | Female | 48 | Caucasian | No | No | no | Myalgia/Headache | G1 | Myalgia | G1 |
| PZB7 | Pfizer Biontech | Female | 34 | Caucasia/Arab | No | Hashimoto thyroiditis | no | None | NA | None | NA |
| PZB8 | Pfizer Biontech | Female | 29 | Arab | No | No | no | Myalgia/Swelling | G1 | Fever/Myalgia | G1 |
| PZB9 | Pfizer Biontech | Male | 41 | Arab | No | Allergic rhinitis | no | Myalgia | G1 | Myalgia | G1 |
| PZB10 | Pfizer Biontech | Female | 35 | Arab | No | No | no | Myalgia | G1 | Fever/Insomnia/Myalgia/Fatigue | G2 |
| PZB11 | Pfizer Biontech | Male | 41 | Caucasian | No | No | no | Myalgia | G2 | Myalgia | G1 |
| PZB12 | Pfizer Biontech | Female | 34 | Indian | No | Hypothyroidism | Levothyroxine | None | NA | Fever/Myalgia | G1 |
| PZB13 | Pfizer Biontech | Male | 29 | Indian | No | No | no | Fatigue | G1 | Fever/Heaviness in arm | G2 |
| PZB14 | Pfizer Biontech | Female | 38 | Arab | No | Allergic | no | Myalgia/Headache | G1 | Fatigue | G1 |
| PZB15 | Pfizer Biontech | Male | 43 | Arab | No | Hypothyroidism | Levothyroxine | None | NA | Myalgia/Headache | G2 |
| PZB16 | Pfizer Biontech | Female | 39 | Indian | No | No | no | Heaviness in arm | G1 | Fever/Myalgia/Fatigue | G2 |
| PZB17 | Pfizer Biontech | Female | 42 | Indian | Yes | T2D, Hypertension | Methformin, Telmisartan | Fever/Headache/Myalgia/Fatigue | G2 | Fatigue/Gastritis | G2 |
| MDA18 | Moderna | Male | 42 | Caucasian | No | Hypertension | Amlodipine, Ramipril | Myalgia | G1 | Myalgia | G1 |
| PZB19 | Pfizer Biontech | Female | 39 | Caucasian | No | No | no | Myalgia | G1 | Headache/Myalgia/Arthralgia | G3 |
| MDA20 | Moderna | Female | 36 | Arab | Yes | No | no | Chills/Myalgia | G2 | Chills/Myalgia | G2 |
| MDA21 | Moderna | Female | 36 | Caucasian | No | No | no | Myalgia | G1 | Fever/Skin rash/Myalgia | G2 |
| PZB25 | Pfizer Biontech | Female | 39 | Caucasian | No | No | no | Myalgia | G1 | Myalgia | G1 |
| MDA26 | Moderna | Female | 30 | Caucasian | Yes | Asthma | Sereti<br>de,<br>Salbut<br>amol | Fever/Head<br>ache/Myalgi<br>a/Fatigue | G3 | Asthma<br>attack/Fever<br>/Myalgia/Fati<br>gue | G3 |

T2D: Type 2 Diabetes
Vaccine type and lot, and subjects' characteristics were recorded, including demographic, biometric data, blood group, underlying diseases, drugs usage, and previous COVID-19 disease.
Every subject recorded and graded the symptoms that occurred after the first and second vaccinations doses, according to the NIH "DAIDS AE Grading Table"

Altogether, the implementation of this protocol established the feasibility of obtaining stabilized-RNA blood samples from study-subjects post-vaccination at high-temporal frequencies. We generated a large dataset using a cost-effective RNA-sequencing protocol that served as the basis for subsequent analyses presented in this paper and was deposited in a public repository. A detailed map of the serological profiles of the subjects enrolled in the study was obtained that permitted us to explore the possible associations between blood transcriptional responses and vaccine immunogenicity.

### The post-prime interferon response peaks at day-2 and correlates with the antibody response

Vaccines can elicit innate immune responses that are detectable systemically via blood transcriptome profiling. But not all of them do, which is for instance the case of the aluminum- adjuvanted Hepatitis B vaccine (9). Therefore our first question was whether transcriptional changes could be observed during the first few days following the administration of COVID- 19 mRNA vaccines.

Analyses were carried out employing a fixed repertoire of 382 transcriptional modules (BloodGen3) that we had recently established and characterized functionally (10)(see methods section for details). Module responses were determined across all time points. The differential gene-set enrichment functions of the dearseq R package were run to assess whether changes observed throughout the nine days post-prime were statistically significant (11). This analysis identified significant temporal changes for 22 of the 382 modules constituting the BlooGen3 repertoire (**Supplementary File 2**).

Only seven modules were found to be changed at any given time point during the first three days following the administration of the priming dose of the vaccine (**Figure 2A**). The abundance of four modules was consistently increased across these time points, and all four belonged to the module aggregate A28. Each “module aggregate” regroups sets of modules that showed consistent abundance profiles across a reference set of 16 disease cohorts that were employed for the construction of the BloodGen3 repertoire (see methods and (10) for details). The module aggregate in question (“Aggregate A28”) comprises of six modules. As described in detail in one of our recent publications, all six are associated with interferon responses (10). The gene composition of the modules and the functional annotations are provided herein (relevant information is provided in **Supplementary File 3** and can be accessed interactively via: https://prezi.com/view/E34MhxE5uKoZLWZ3KXjG/). The responses observed on days 1 and 2 post-prime were mapped onto fingerprint grid plots, where modules occupy a fixed position and are arranged by aggregate. Each aggregate occupyies a given row (**Figure 2A**). Time-course gene-set enrichment analysis confirmed that changes observed over time in four out of six A28 modules were significant. The response profiles of the A28 modules showed a peak on day 2 post-vaccination. This was also visible on a heatmap showing responses at each timepoint across individual subjects (**Figure 2B**). We next examined whether this signature correlated with antibody responses measured 14-days post-prime as well as at 14-days post-boost (**Figure 2C**). For this, correlation analyses were run at the module level within Aggregate A28 using, as the endpoint, fold-changes in antibody levels on days 7 and 14 post-prime and days 7 and 14 post-boost relative to the pre- vaccination baseline (immediately prior to the administration of the first dose of COVID-19 mRNA vaccines). “Significance hotspots” were identified when most modules within a given aggregate reached correlation significance thresholds. In the case of the post-prime interferon signature, we identified such significance hotspots on days 2 and 3 post-prime for a subset of three interferon modules, M10.1, M15.127, and M83, while a fourth module, M15.86, also displayed significant correlations across all antibody types, but only on day 2.

**Figure 2:**
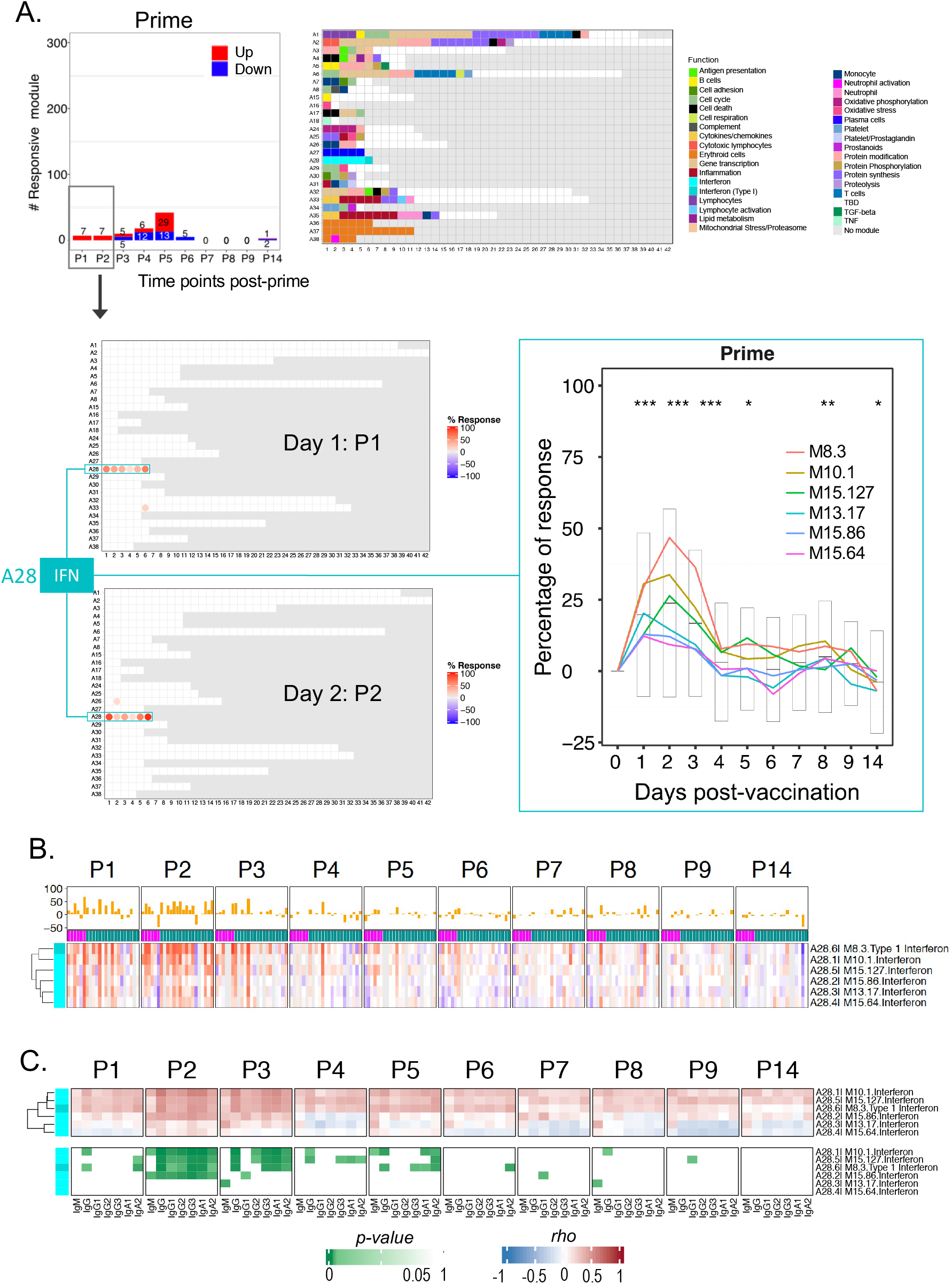
Characterization of the post-prime interferon response signature. (A) The bar graph shows the cumulative module response at the various timepoints following the administration of the priming dose of the vaccine (noted P1-P14). The Y-axis values and numbers on the bars indicate the number of modules meeting the 15% response threshold (out of a total of 382 modules constituting the BloodGen3 repertoire, with percentage response corresponding to the proportion of transcripts predominantly increased or decreased compared to baseline using FDR < 0.1 as the cutoff to determine significance [DESeq2]). The number of modules for which abundance was predominantly increased is shown in red, and those for which abundance was predominantly decreased are shown in blue. The fingerprint grid plots represent the overall module responses on day 1 post-prime (P1) and day 2 post- prime (P2). Modules from the BloodGen3 repertoire occupy fixed positions on the fingerprint grids. They are arranged as rows based on membership to module aggregates (rows A1 through A38). Changes compared to the pre-vaccination baseline are indicated on the grid by red and blue spots of varying color intensity, which indicate the “percentage response” for a given module. The color key at the top indicates the various functions attributed to the modules that are represented on the grid. The response of the six modules comprising aggregate A28 is represented on a line graph that shows the proportion of responsive transcripts for each module across all the post- prime timepoints. For each module, the statistical significance of the overall response was determined by time- course gene set enrichment analysis. Four of the six A28 modules met significance thresholds FDR < 0.1 (M8.3: p- value = 1.9-e4, FDR = 0.019, M10.1: p-value = 1.9-e4, FDR = 0.019, M15.127: p-value = 1.9-e4, FDR = 0.019, 727 and M15.86: p-value = 3.9-e4, FDR = 0.031) and all six A28 modules p < 0.05 (M13.17: p-value = 1.5-e3, FDR = 0.101 and M15.64: p-value = 0.044, FDR = 0.727). We also ascertained the significance of changes measured post-prime at the level of this module aggregate and at each time point (paired t-test comparing module response at each time point relative to the pre-vaccination baseline; * p<0.01, ** p<0.001, *** p<0.0001). (B) Heatmaps represent proportions of transcripts that changed within the six A28 modules at different timepoints and across different individuals compared to pre-vaccination baseline values. Red indicates that transcripts were predominantly increased over the baseline, and blue indicates that transcripts were predominantly decreased. Rows represent the six A28 modules arranged within an aggregate via hierarchical clustering. Columns represent samples grouped by timepoint and show profiles of individual subjects within each timepoint. (C) The heatmaps represent associations (Spearman correlation test) between antibody responses measured 14 days after administration of COVID-19 booster doses and transcriptional responses measured across nine consecutive days after the priming dose. The heatmap at the top provides the correlation coefficients across multiple days and for each day across multiple subjects, with rows corresponding to the six A28 interferon modules. The heatmap below shows the significance of the correlations shown on the heatmap directly above, with the same ordering of rows and columns.

Thus, we found that an interferon response is induced over the first three days following the administration of the priming dose of mRNA vaccines. Remarkably this signature correlated with the antibody response measured several weeks later, 14 days after the administration of the second dose of vaccine.

### A decrease in inflammation is accompanied by an increase in adaptive immune response genes on day 5 post-prime

We were next interested in characterizing the changes occurring beyond the first three days following administration of the priming dose. In total, 18 modules displayed changes on day 4 post-prime, of which 12 showed a decrease in abundance. These modules belonged to three aggregates that have been associated with inflammation (A31, A33, A35). Most changes were observed on day 4, but for some modules, changes were apparent starting on day 3 and continued beyond days 4, day 5, or even 6 (**Supplementary Figure 1**). In our earlier work, modules within the BloodGen3 Aggregate A35 were associated with systemic inflammation mediated by neutrophils and were found to constitute a common denominator across a wide range of pathologies in which systemic inflammation is present (12). The association of A35 with inflammatory processes was also ascertained based on the results of the functional profiling analyses and the restriction in transcript expression in the reference datasets (10). Detailed functional annotations can be accessed via interactive circle packing charts: https://prezi.com/view/7Q20FyW6Hrs5NjMaTUyW/). Module Aggregate A33 has not been investigated as extensively in any of our prior studies but was clearly associated with inflammation via functional profiling (https://prezi.com/view/VBqKqHuLWCra3OJOIZRR/).

The peak response post prime was on day 5, with a total of 42 modules showing differences in comparison to the pre-vaccination baseline (**Figure 3**). At this timepoint, most modules showed an increase in abundance (29 were increased and 13 decreased). Some of those modules belonged to aggregates that were associated with adaptive immunity, most notably A27, which is associated with plasmablast responses (three out of five modules were responsive at this timepoint). This association is based on the restriction of the expression of the genes comprising A27 modules in plasma cells observed in a reference dataset including a wide range of cell populations (contributed by Monaco et al. (14)) and by the presence of the plasmablast marker CD38 and other associated genes (*IGJ*, *TNFRSF17*, *TXNDC5*) in one of the A27 modules (M12.15). Detailed annotations and expression profiles of A27 transcripts in the reference datasets can be accessed via https://prezi.com/view/GgIiA0K9kSFHbpVj2I85/. Other immune-relevant modules found to be increased at this timepoint are associated with T-cells (M12.6 from aggregate A1). Detailed functional annotations for module aggregate A1 can be accessed via https://prezi.com/view/sxap39tKxkmCNTTNIlVO/). Others were mapped to module aggregates A24 and were associated with oxidative phosphorylation which is known to play a role for instance in T-cell activation (6 out of 11 modules were responsive) (13). Other modules were not yet functionally annotated, including for instance the four responsive modules, out of 15, belonging to aggregate A26. Notably, the signatures observed on day 5 appeared to be transient, and no modules were increased on day 6 post-prime.

**Figure 3:**
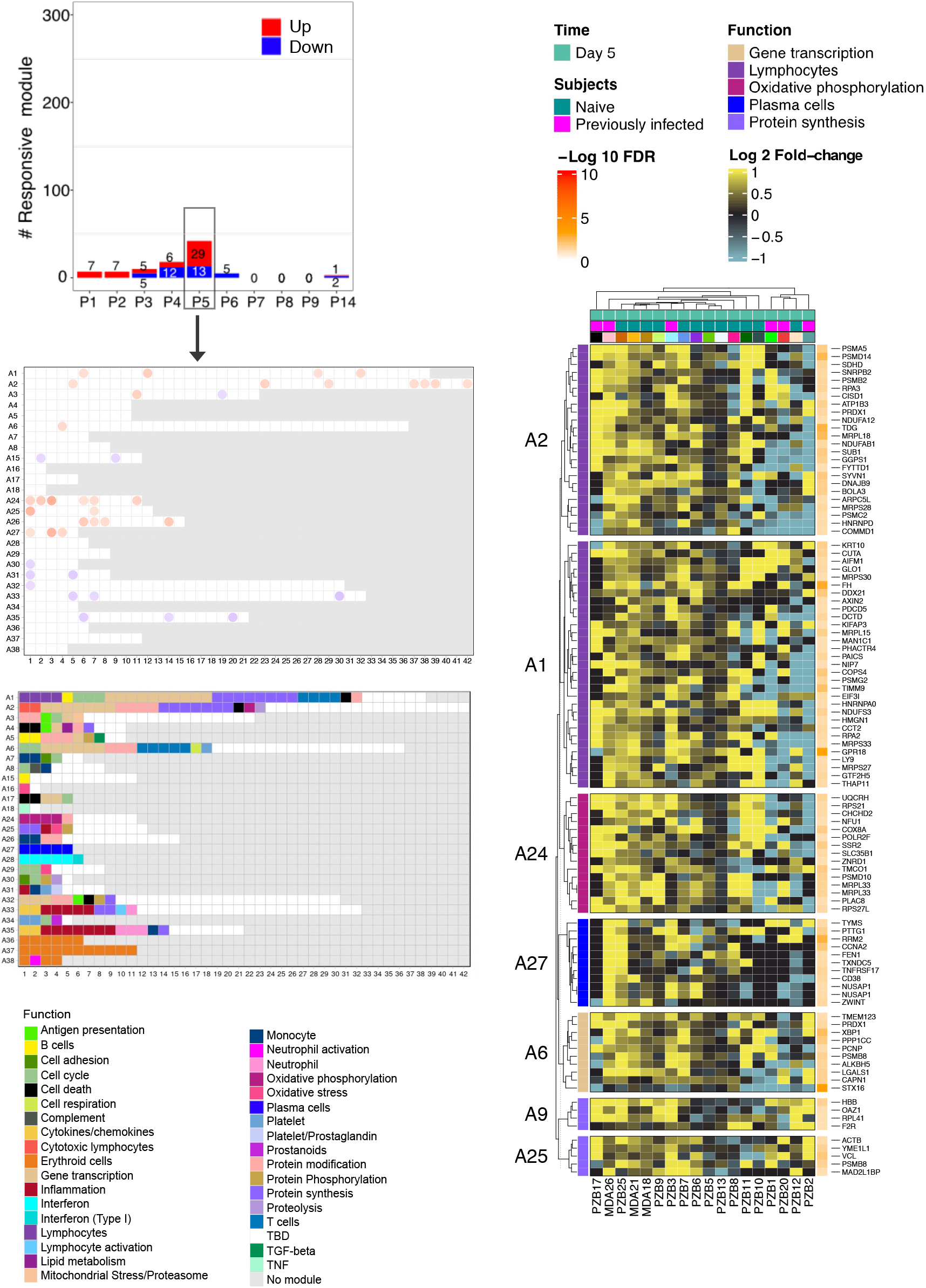
Characterization of responses on day 5 post-prime. **(A)** The bar graph shows the cumulative number of responsive modules at each timepoint following the administration of the priming dose of the vaccine (noted P1- P14). The fingerprint grid plot shows changes observed at P5 (day 5 post-prime). The position of the modules on the grid is fixed. The percent response of individual modules is represented on the grid by red and blue spots of varying color intensity denoting a predominant increase or decrease in abundance, respectively. The percentage response of a given module corresponds to the proportion of transcripts predominantly increased or decreased compared to baseline, meeting a significance cutoff of FDR < 0.1. The color key at the top indicates the various functions attributed to the modules that are represented on the grid. **(B)** The heatmap represents Log2 average fold change in abundance of transcripts constituting sets of modules associated with given functional annotations on P5. Rows represent individual transcripts grouped according to the module aggregate they originate from, corresponding to the different rows on the fingerprint grid plot on the left. Each module aggregate is associated with a unique function, as indicated by the color key above. The columns on the heatmap represent individual subjects coded with the type of vaccine received (Pfizer BioNtech = PZB; Moderna = MDA).

Taken together, we found the number of responsive modules to peak on day 5 post-prime. A decrease in the abundance of transcripts associated with inflammation was accompanied by an increase in the abundance of transcripts associated with adaptive immune responses. Notably, the latter appeared earlier than seen in response to other vaccines where plasmablast signatures are observed around day 7 post-vaccination (6,14,15).

### A post-boost interferon signature peaks on day 1 and correlates with antibody responses

After delineating temporal responses post-prime, we examined changes after the second dose of COVID-19 mRNA vaccines. Time-course gene set enrichment analysis identified significant temporal changes for 311 of 382 modules comprising the BloodGen3 repertoire (**Supplementary File 4**). After the booster dose, the peak number of responsive modules occurred on day 1, with 261 responsive modules or about two-thirds of the 382 modules constituting the BloodGen3 repertoire (**Figure 4**). This number decreased sharply afterward, with 115 responsive modules on day 2 and only 9 responsive modules on day 3. The kinetic and amplitude of the post-boost response contrasted markedly with that observed post-prime, when, as described above, the number of responsive modules after the first dose instead peaked on day 5, with changes found in 42 modules at that timepoint.

**Figure 4:**
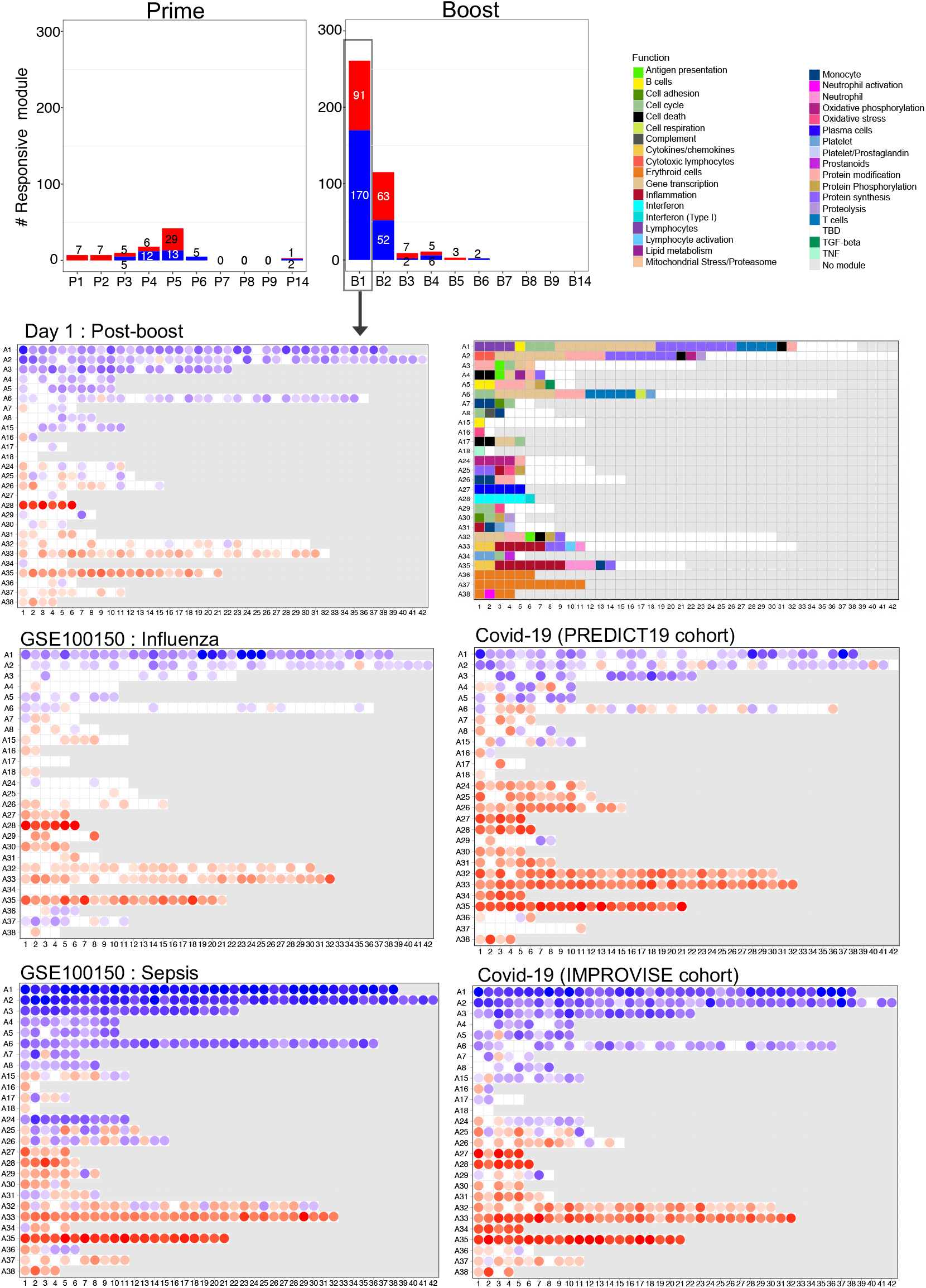
Fingerprint grid plots mapping changes observed on day 1 post-boost and across reference datasets. The bar graphs show the cumulative module response at the various timepoints post-priming and booster doses (noted P1-P14 and B1-B14, respectively). The Y-axis values and numbers on the bars indicate the number of modules meeting the 15% response threshold (out of a total of 382 modules constituting the BloodGen3 repertoire, with percentage response corresponding to the proportion of transcripts predominantly increased or decreased compared to baseline meeting a significance cutoff of DESeq2, FDR < 0.1. The fingerprint grid plots show changes in transcript abundance in a given study group in comparison to baseline (pre-vaccination sample or uninfected control group – with the percent response of individual modules shown by red and blue spots of varying color intensity denoting predominant increase or decrease in abundance, respectively. Changes are shown in the top grid for a group comparison of 1 day after receiving the booster dose of COVID-19 mRNA vaccines with baseline pre-vaccination samples (this study). Grids in the middle and bottom positions show changes for patients with acute infections caused by influenza virus (public dataset) or SARS-CoV-2 (this study) and for patients with bacterial sepsis (public dataset). The color key at the top indicates the various functions attributed to the modules that occupy a fixed position on the grid plot.

As seen from the fingerprint grid plot, the day 1 post-boost response was extensive and polyfunctional (**Figure 4**). An overall decrease in abundance was observed for aggregates broadly associated with lymphocytic cells (Aggregates A1-A8) and increased for module aggregates associated with myeloid cells, inflammation, and circulating erythroid cells (Aggregates A33-A38). In addition, a marked increase in the abundance of modules associated with interferon responses was also observed (Aggregate A28). We compared the day 1 response fingerprint of the COVID-19 mRNA booster vaccine to fingerprints derived from patients with a wide range of pathologies. These included sixteen reference datasets encompassing infectious and autoimmune diseases, as well as cancer, solid organ transplant recipients, among others (these cohorts are described in our previously published work (10, 16); the respective blood transcriptome fingerprint collections are accessible via a dedicated web application: https://drinchai.shinyapps.io/BloodGen3Module/). In addition, we analyzed two original COVID-19 blood transcriptome datasets: one cohort comprising 77

Covid-19 patients with disease severities ranging from mild and moderate to severe (the “PREDICT-19 consortium Italian cohort dataset” – see methods and published study protocol for details (17)), while the second cohort comprised 40 COVID-19 patients recruited at the time of admission to the intensive care unit (ICU) (“IMPROVISE cohort whole blood dataset”). These high-level comparisons showed, firstly, that the extent of the changes associated with the day 1 response to the second dose of the COVID-19 mRNA vaccine was consistent with that observed in some patient cohorts with acute infections (**Figure 4**). More specifically, they were found to most resemble the responses seen in a cohort of subjects with influenza infection, with a marked interferon response (A28) and an inflammation signature (A33, A35). At a higher level, these response patterns were also generally consistent with those observed in patients with a COVID-19 infection. However, the changes that occurred in response to the vaccinations were not as extreme as those found, for instance, in patients with sepsis or with the most severe form of COVID-19 (i.e., the IMPROVISE dataset) (most notably for inflammation [A33, A35] and erythroid cell responses [A36-A38]).

Overall, the BloodGen3 transcriptome fingerprint observed on day 1 after the second vaccine dose contrasted markedly with the fingerprint observed on day 1 post-prime. Yet, the interferon response signature was found to be a common denominator between the responses to the first and second doses, as it was observed in both cases in the first few days following administration of the vaccine. We therefore began to dissect the post-boost response by examining this interferon response signature in more detail.

Following the administration of the booster dose, the interferon response was noticeably sharper in comparison to the post-prime response and peaked on day 1 instead of day 2 (**Figure 5A**). This was illustrated by the difference in the maximum average module response, which was close to 50% of the constitutive transcripts on day 2 post-prime and greater than 80% on day 1 post-boost.

**Figure 5:**
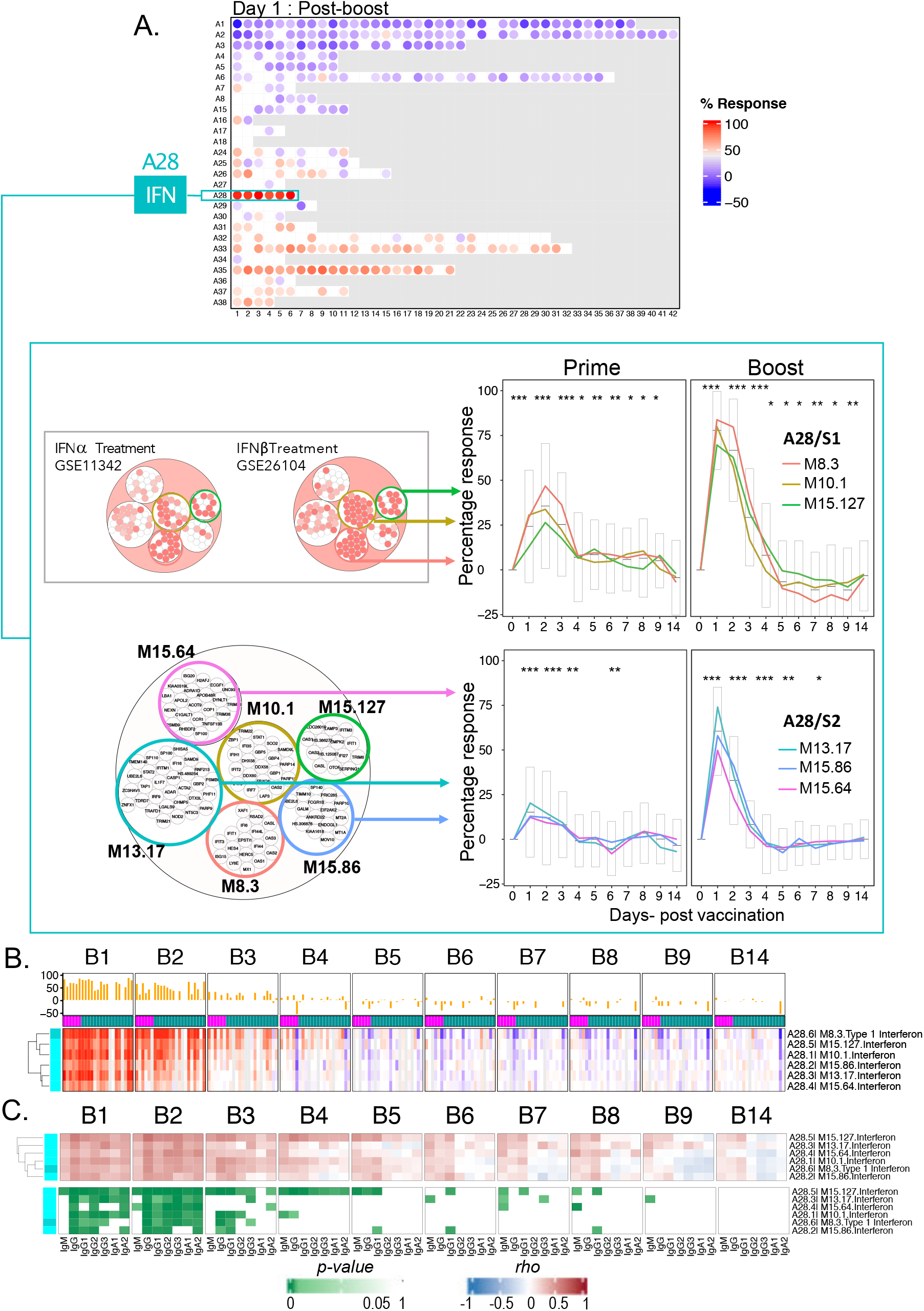
Characterization of the day 1 post-boost interferon response signature. **(A)** The fingerprint grid plot maps the modular response observed on day 1 post-boost (percent response is determined based on statistical cutoff: DESeq2, FRD < 0.1). The six modules forming the A28 aggregate are highlighted. The line graphs below represent the summarized percentage responses at the module level, encompassing all study subjects (one line per module). Percentage response accounts for the proportion of transcripts for a given module showing differences in abundance post-prime (left) or post-boost (right) compared to baseline pre-vaccination levels. Changes in transcript abundance post-prime and post-boost for two distinct sets of interferon response modules that received the denomination A28/S1 and A28/S2 are plotted on separate graphs. For each module, statistical significance for the overall response was determined by time course gene set enrichment analysis. Significance was reported post- prime in Figure 2. Post-boost all six A28 modules met significance thresholds p < 0.001 and FDR < 0.001 (M8.3: p-value = 1.9-e4, FDR = 3.6-e4, M10.1: p-value = 1.9-e4, FDR = 3.6-e4, M13.17: p-value = 1.9-e4, FDR = 3.6-e4, M15.127: p-value = 1.9-e4, FDR = 3.6-e4, M15.64: p-value = 1.9-e4, FDR = 3.6-e4 and M15.86: p-value = 1.9-e4, FDR = 3.6-e4). In addition, we ascertained the significance of changes measured post-prime at the level of this module aggregate and at each time point (paired t-test comparing module response at each time point relative to the pre- vaccination baseline; * p<0.01, ** p<0.001, *** p<0.0001). The circle packing plots on the left show module responses at the individual transcript level for two public blood transcriptome datasets. The larger circle below indicates official symbols for the individual transcripts. It also highlights the modules included in A28/S2, shown directly on the right. The smaller circles above show changes in abundance of A28 transcripts for two public datasets. One study (GSE11342) measured blood transcriptional response in patients with Hepatitis C infection treated with alpha-interferon (23). The second study (GSE26104) measured transcriptional response in subjects with multiple sclerosis treated with beta-interferon (24). A red circle indicates a significant increase in the abundance of transcripts compared to the pre-treatment baseline (*|fold-change| > 1.5, FDR < 0.1). **(B)** Changes in abundance compared to baseline pre-vaccination levels are represented on a heatmap, with modules as rows and individual samples as columns. The modules are arranged by hierarchical clustering based on abundance patterns across samples. The samples are arranged by timepoints post-prime (top) and post-boost (bottom). **(C)** The heatmaps represent associations (Spearman correlation) between antibody responses measured 14 days after administration of COVID-19 booster doses and transcriptional responses measured across nine consecutive days after the booster dose. The heatmap on top provides the correlation coefficients across multiple days and for each day across multiple subjects, with rows corresponding to the six A28 interferon modules. The heatmap below shows the significance of the correlations shown on the heatmap on top, with the same ordering of rows and columns.

We decided to then perform hierarchical clustering to identify subsets of modules within the A28 aggregates that might group together based on patterns of transcript abundance across all subjects and timepoints. Two sets of three modules each were, thus, identified within the A28 aggregate. The first set comprised modules M8.3, M10.1, and M15.127 (referred to as A28/S1), and the second set comprised modules M16.64, M13.17, and M15.86 (referred to as A28/S2). Interestingly, we observed post-prime that, while modules in A28/S1 peaked on day 2, those belonging to A28/S2 peaked on day 1 (**Figure 5B**). Furthermore, A28/S1 modules showed an extended peak post-boost, with day 2 levels being almost identical to those of the day 1 peak, while A28/S1 modules peaked sharply on day 1, with levels decreasing rapidly thereafter. These findings suggest that both sets of modules measured distinct types of interferon response. Indeed, public datasets in which responses to type 1 interferon were measured *in-vivo* indicated that A28/S1 modules are likely to represent type 1 interferon responses (**Figure 5B**), while we postulated that A28/S2 modules might represent a type 2 interferon response. Modules forming the A28/S1 set comprise some of the better recognized “canonical” interferon response genes, such as Oligoadenylate Synthetase family members (OAS1, OAS2, OAS3, OASL), Interferon Induced Protein family members (IFI6, IFI27, IFI35, IFI44, IFI44L), as well as Interferon Induced Protein With Tetratricopeptide Repeats family members (IFIT1, IFIT3, IFIT5) (10). Modules forming the A28/S2 set comprise instead most notably members of the Nuclear Antigen family members SP100, SP110 and SP140, which are associated with interferon gamma signaling, as well as transcription factors IRF9 and STAT2. Composition and functional annotations for A28 modules can be explored further at: https://prezi.com/view/E34MhxE5uKoZLWZ3KXjG/.

Finally, a strong association was found between the post-boost interferon signature and the subsequent development of an antibody response. Indeed, positive correlations were observed for all six A28 modules that reached significance on days 1, 2, and 3 post-boost. Notably, this differed from the post-prime interferon response, for which significance was reached only for four of the six modules and only on days 2 and 3.

Taken together, the high temporal resolution profiling results permitted the delineation of distinct patterns of post-prime and post-boost interferon responses. The timing of the responses observed at the individual module level contributed to the definition of the two distinct sets of interferon modules. One set was associated with responses to type I interferon *in-vivo* and dominated the post-prime response, with a peak on day 2. The post-boost response showed a strong induction of both sets and also peaked on day 1.

### Inflammation and erythroid cell signatures peak sharply on day 1 post-boost

We continued the dissection of the day 1 post-boost signature, focusing this time on responses associated with inflammation and circulating erythroid cell precursors.

Aggregates A33 and A35, which are associated with inflammation, tended to decrease from day 4 through day 6 post-prime but displayed instead a sharp and transient increase in abundance post-boost. Indeed, a well-delineated response peak was observed on day 1 post-boost for both the A33 and A35 modules (**Figure 6**), but in contrast to the interferon response (A28/S1), it did not extend beyond the first day. Three distinct response patterns were identified via hierarchical clustering among the 21 modules that formed aggregate A35. The “A35/S1” set comprised five modules, while “A35/S2” and “A35/S3” comprised ten and six modules, respectively. The distinction between those three A35/inflammation module sets was rather more subtle than was the case for the A28/interferon sets. Indeed, all three module sets peaked on day 1 post-boost. Differences were rather apparent in the inflection of changes measured on days 2 and 3 post-boost and in the “recovery phase”, as abundances appeared to dip below the baseline and progressively rise to reach pre-vaccination levels. The underlying biological factors driving the grouping of the modules to those three distinct sets could not be identified at this time.

**Figure 6:**
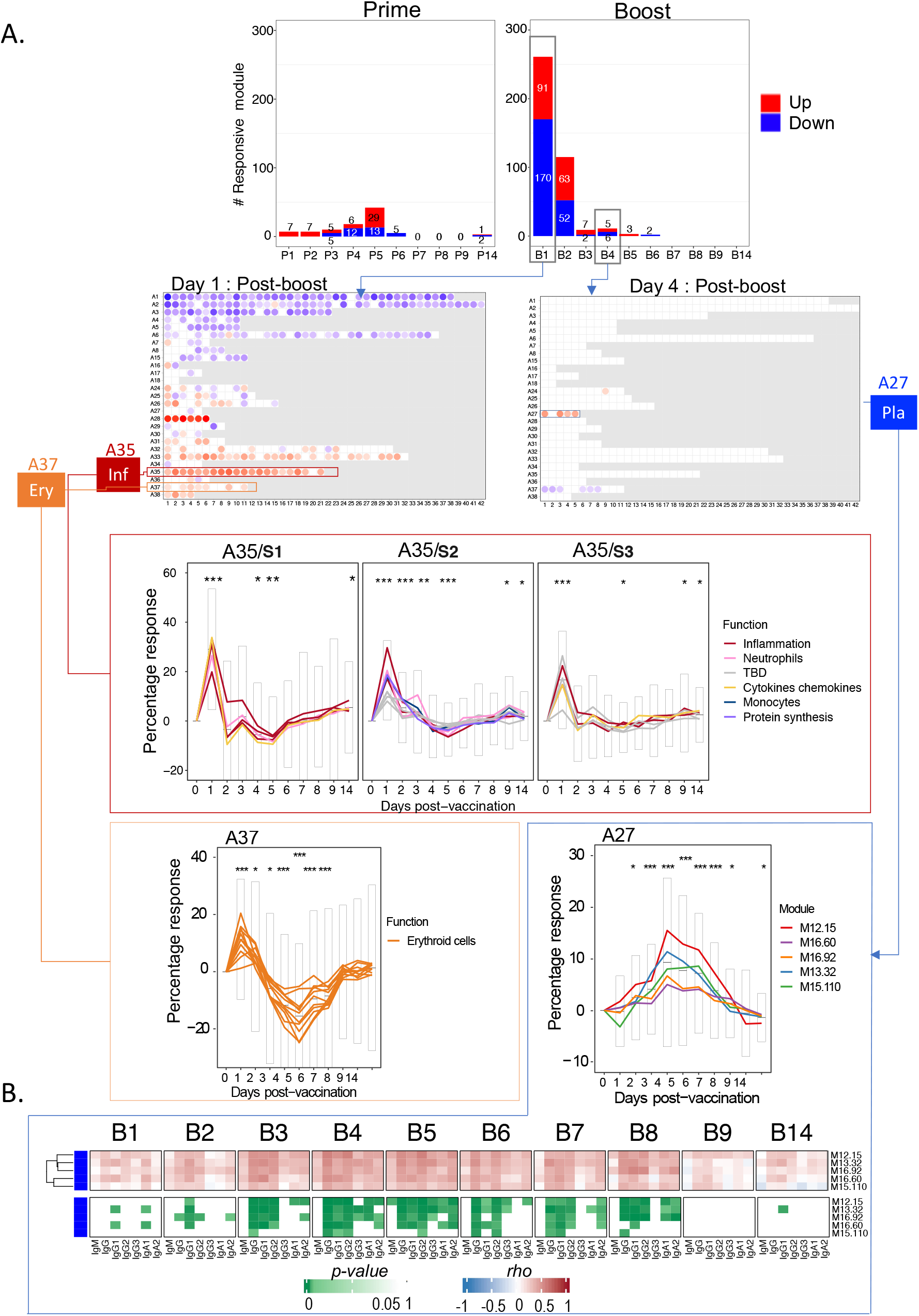
Characterization of post-boost inflammation, erythroid cell, and plasmablast responses. **(A)** The bar graph at the top represents the number of response modules at any given time point post-prime and post-boost. The fingerprint grid plots show the modules that had changes compared with a fixed visualization and interpretation framework. Changes are shown for the day 1 post-boost timepoint (left) as well as day 5 (right) (percent response is determined based on statistical cutoff: DESeq2, FRD < 0.1). On the left grid, modules belonging to aggregates A35 (associated with inflammation) and A37 (associated with erythroid cells) are highlighted. The profiles of those modules are represented on the line graphs below, which show the average percentage responses of A35 and A37 modules across multiple timepoints. The percentage response for a given module is the proportion of its constitutive transcripts showing significant changes, ranging from 0% to 100% when transcripts were predominantly increased to 0% to −100% when transcripts were predominantly decreased. Each line represents the profile of the modules constituting a given aggregate. Line graphs for A35 were split into three sets according to differences in clustering patterns (A35/S1, A35/S2, and A35/S3). On the right grid, modules belonging to aggregates A27 (associated with platelets) are highlighted. The corresponding line graph below represents the changes in abundance of A27 modules over time following administration of the second dose of vaccine. For each module, statistical significance for the overall response was determined by time course gene set enrichment analysis using the dearseq R package. For A35, 20 of 21 modules met significance thresholds (p-value < 0.05 and FDR < 0.01). It was also the case in 11 of 11 modules for A37 and 4 of 5 modules for A27 (**Supplementary file 4**). In addition, we ascertained the significance of changes measured post-prime at the level of this module aggregate and at each time point (paired t-test comparing module response at each time point relative to the pre-vaccination baseline; * p<0.01, ** p<0.001, *** p<0.0001). **(B).** The heatmaps represent associations between antibody responses measured 14 days after administration of COVID-19 booster doses and transcriptional responses measured across nine consecutive days after the booster dose. Specifically, the heatmap at the top represents the correlation coefficients across multiple days and for each day across multiple subjects, with rows corresponding to the five A27 plasmablast modules. The heatmap below shows the significance of the correlations shown on the heatmap at the top, with the same order of rows and columns.

Modules for three aggregates broadly associated with erythroid cell signatures also displayed a sharp but transient increase in transcript abundance on day 1 post-boost. However, the abundance tended to dip afterward, with a low peak on day 4 post-boost, before recovering by day 7. Functionally, this signature was found to be most prominently associated with immunosuppressive states, such as late-stage cancer or pharmacological immunosuppression (16), which is consistent with published functional studies (18, 19). We also found such signatures were associated with more severe manifestations in babies infected with Respiratory Syncytial Virus (RSV) (16). Moreover, erythroid precursors have been recently associated with COVID most severe clinical outcomes (20). Finally, we did not find evidence of an association between the day 1 post-boost inflammation or erythroid cell signatures and the antibody responses.

### A plasmablast signature peaks on day 4 post-administration of the booster dose and correlates with antibody responses

After the booster dose, the number of responsive modules peaked sharply on day 1, then rapidly subsided beyond day 2, with the number of responsive modules on days 3, 4, 5, and 6 being reduced to 8, 11, 3, and 2, respectively. Yet, changes within this later timeframe are meaningful, as they specifically concern the set of five modules comprising aggregate A27, which is associated with the presence of antibody-producing cells in the peripheral blood.

Three of the five A27 modules showed significant alterations after the booster dose (M16.60, M13.32, M12.15) (**Figure 6**). The proportion of differentially expressed transcripts in each module was relatively modest (with an average of 15% at the peak of response), especially in comparison with the interferon signatures described above (with an average of >80% for some modules at the response peak). Yet, the trajectories of the five A27 modules were relatively consistent, with only one of the modules (M15.110) showing a different pattern, i.e., a peak on day 6, slightly above the levels observed on day 4. We also examined the association of this post-boost plasmablast signature with the antibody response and found a significant association starting from about day 3 and lasting until day 7 post-boost (**Figure 6**). In summary, COVID-19 mRNA vaccination induced a plasmablast response that peaked on day 4 post-vaccination. This was unexpected since such signatures typically are measured around day 7 post-vaccine administration (e.g., in the case of influenza or pneumococcal vaccines (6)). We were also able to demonstrate a logical association between this post-boost plasmablast signature and the subsequent development of humoral immunity.

### Patterns of interferon induction elicited by COVID-19 mRNA vaccines are also observed among COVID-19 patients

Our work identified the interferon response as the most upstream factor associated with the development of humoral immunity following COVID-19 mRNA vaccination. High-temporal resolution profiling identified distinct patterns of interferon induction post-prime and post-boost and we next decided to determine whether similar response patterns could be identified among patients with COVID-19 disease.

We relied for this on the original blood transcriptome data from the PREDICT-19 consortium Italian COVID-19 cohort comprising 77 patients with a wide spectrum of disease severity. We used the response values for the six interferon modules from Aggregate A28 to map individual COVID-19 patient samples along with post-vaccine samples on the same t-SNE plot (**Figure 7A**). First, we confirmed that there was no apparent separation of the vaccination and COVID-19 patient cohorts, and that batch correction was therefore not warranted before proceeding with comparative analyses (**Supplementary Figure 2**). This is consistent with the results of meta-analyses we have previously conducted at the module level (16). To help with the interpretation, k-means clustering was performed using the consolidated set of samples, resulting in the formation of eight distinct clusters. Next, we examined the distribution of samples from the vaccine and COVID-19 cohorts across the tSNE plot and among the eight clusters. Timepoints at which an interferon response was detectable in vaccinated subjects were of particular interest. Indeed, day 1 and day 2 post-prime samples (P1, P2), while preferentially found in Clusters 1 and 5, appeared to be distributed across the entire t-SNE plot. This is in contrast with day 1 and day 2 post-boost vaccination samples (B1, B2), which were almost exclusively found in Cluster 5. A set of COVID-19 patients also co-localized in Cluster 5, while others were found scattered across clusters, especially Clusters 1, 2, 6, and 3. Interferon responses were detectable in all these clusters, but with important nuances. For one, samples from Cluster 5 showed by far the most potent responses, with responses seen in most cases across all six interferon modules, which was consistent with the post-boost vaccine response (**Figure 7B**). In comparison, the response was less pronounced in samples from Cluster 1, which was dominated by modules associated with type I interferon responses (the A28/S1 set comprising M10.1, M8.3 and M15.127 described above). This pattern of response was more consistent with the post-prime vaccine response. Signatures for samples forming Clusters 2 and 6 were not well-defined and were in some cases absent, yet these clusters also included COVID-19 patients. Samples forming Cluster 3 displayed a peculiar signature, with an increase in the abundance of modules belonging to the A28/S2 set (M15.64, M13.17, and M15.86) concomitantly with a decrease in modules forming A28/S1. Among the samples forming this cluster, this pattern was most apparent for the COVID-19 patients.

**Figure 7:**
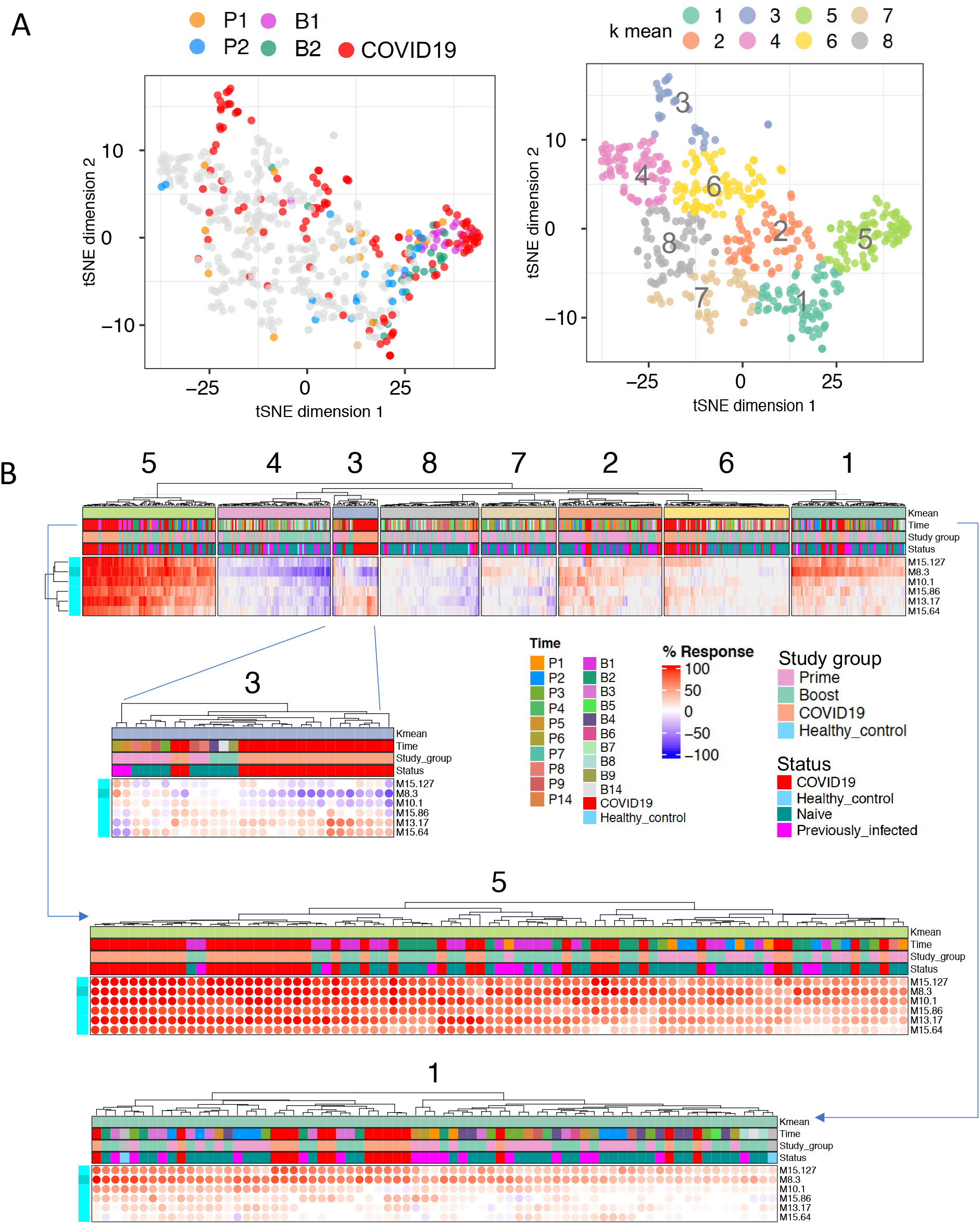
Comparing patterns of interferon response in vaccinated individuals and a cohort of COVID-19 patients. **A.** The tSNE plot represents similarities in patterns of interferon response induction across the six modules forming aggregate A28 and among samples comprised in our vaccination cohort and one of our COVID- 19 disease cohorts (PREDICT-19 / Italy). COVID-19 samples are shown in red along with specific post-vaccination timepoints (post-prime days 1 and 2 [P1, P2], post-boost days 1 and 2 [B1, B2]). Samples from the consolidated cohorts were partitioned into 8 clusters via k-means clustering, the distribution of which is shown on the tSNE plot on the top right. **B**. Heatmaps show patterns of response for the six interferon response modules across the eight sample clusters. The red colors indicate that the abundance of transcripts for a given module is predominantly increased with the intensity representing the proportion of constitutive transcripts meeting a given threshold, which at the level of individual samples is a fixed fold change and difference cutoff (|Fold change| > 1.5, and |difference| > 10 in a given sample over its respective pre-vaccination baseline). The blue color denotes a predominant decrease in abundance of constitutive transcripts compared to the same individual’s pre-vaccination

Thus, we employed here the interferon responses observed post COVID-19 vaccination as a benchmark for the interpretation of COVID-19 patient signature. We were able to establish that most COVID-19 patients display responses consistent with those found post-vaccination, which, as established in this study, were associated with the development of potent humoral responses. However, a subset of patients displayed patterns of interferon response that are not typically seen in vaccinated individuals. It can thus be surmised that the later patterns of interferon response might either be suboptimal or possibly even pathogenic.

### The atypical interferon response signature observed in COVID-19 patients is associated with a worse course of disease

The fact that some COVID-19 patients failed to display robust “post-vaccine-like” interferon responses may be due to either a defective innate immune response, which may lead to more severe disease course, or conversely to activation thresholds not being reached in patients presented with milder disease.

Thus, we next examined patterns of interferon response in another original COVID-19 disease cohort, comprised exclusively of patients enrolled at the time of admission in the ICU (the IMPROVISE cohort, which was also described above). As described above, we again mapped individual COVID-19 patient samples along with post-vaccine samples on a t-SNE plot based on similarities in the patterns of interferon responsiveness across the six A28 interferon modules (**Figure 8A**). COVID-19 subjects were found to again be distributed throughout multiple clusters. Patients who co-localized with day 1 post-boost vaccine samples tended to have relatively short ICU stays (in Cluster 5 with potent A28/S1 and A28/S2 responses), and only a few patients co-localized with day 2 post-prime samples in Cluster 3, which was characterized by a more prominent A28/S1 signature compared with A28/S2. Furthermore, distinct groups of patients in Clusters 1 and 6 displayed the peculiar pattern of interferon response dominated by A28/S2 that was identified earlier among patients enrolled in the PREDICT-19 cohort. Notably, patients from the IMPROVISE cohort displaying this pattern of interferon response showed significantly lengthier stays in the ICU compared to patients displaying patterns of interferon response that are consistent with those observed post-vaccination (**Figure 8B** comparing left and right cluster: for length of hospital stay, t-test, p = 0.006 (**), mechanical ventilation days p =0.016 (*) and ICU stay p = 0.012(*)).

**Figure 8:**
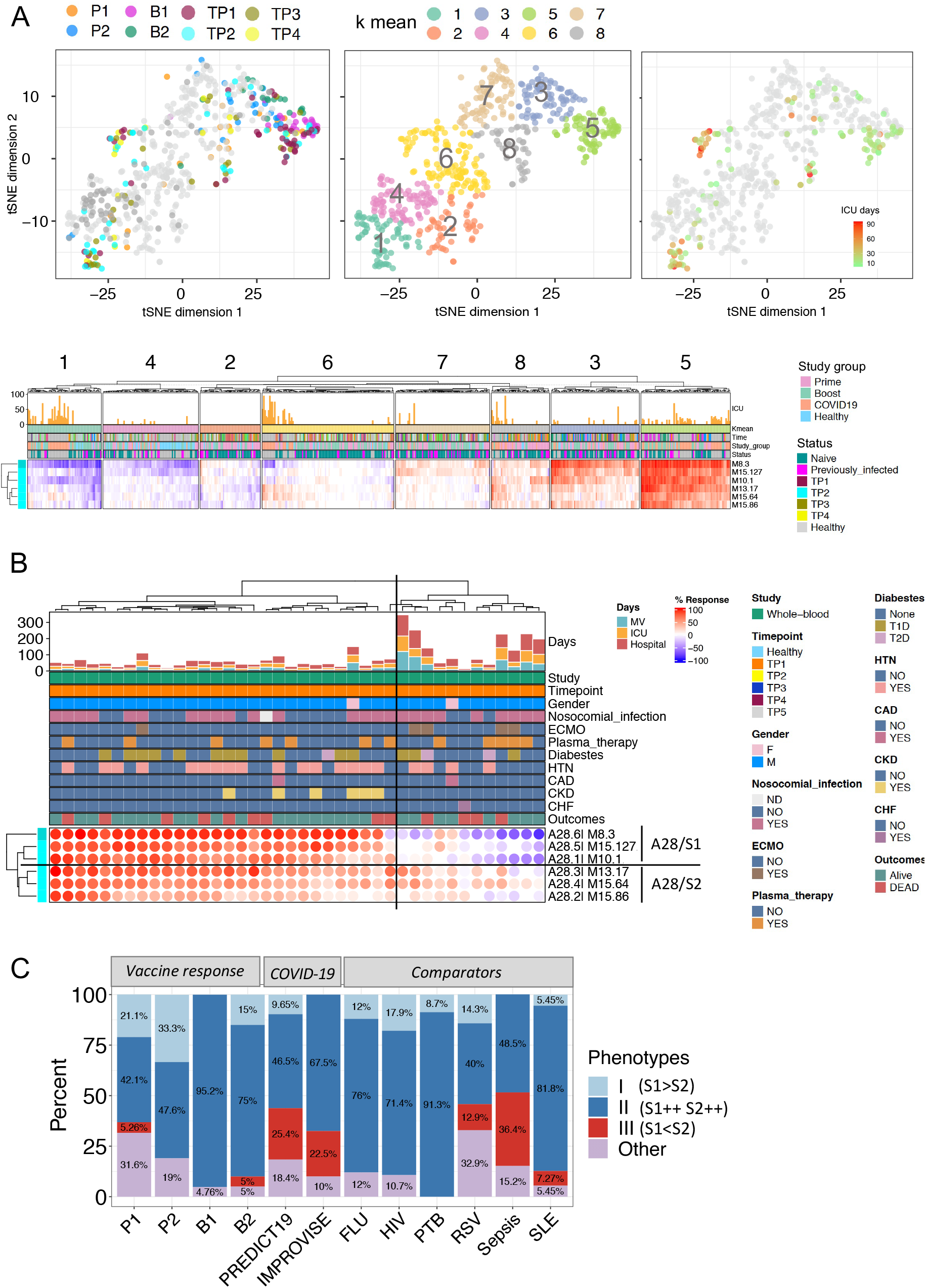
Comparison of interferon response patterns of vaccinated individuals and a cohort of COVID-19 patients with severe disease under intensive care. **A.** The tSNE plot represents similarities in patterns of interferon response induction across the six modules forming aggregate A28 and among samples comprised in our vaccination cohort and one of our COVID-19 disease cohorts (IMPROVISE). Specific post-vaccination timepoints (post-prime days 1 and 2 [P1, P2], post-boost days 1 and 2 [B1, B2]), as well as repeat sampling for COVID-19 patient (TP1, TP2, TP3, TP4, all collected during ICU stay) are shown on the plot on the left. Samples from the consolidated cohorts were partitioned into 8 clusters via k-means clustering, the distribution of which is shown on the tSNE plot on the center. Length of ICU stay is shown on the tSNE plot on the right. Patterns of response for the six interferon response modules across the eight sample clusters are shown on a heatmap below. The red colors indicate that the abundance of transcripts for a given module is predominantly increased with the intensity representing the fold change and difference cutoff (|Fold change |> 1.5, and |difference| > 10 in a given sample over its respective pre-vaccination baseline). The blue color denotes a predominant decrease in abundance of constitutive transcripts compared to the same individual’s pre-vaccination baseline. **B.** The heatmap shows patterns of interferon responses for COVID-19 patients with severe disease upon ICU admission. Multiple clinical parameters are shown on the tracks above (ECMO [Extracorporeal Membrane Oxygenation], HTN [Hypertension], CAD [Coronary Artery Disease], CKD [Chronic Kidney Disease], CHF [Congestive Heart Failure]). The histogram represents the length of stay in the hospital, in the ICU, and under mechanical ventilation, in days. **C.** The bar graph represents for different datasets the proportion of samples corresponding to Interferon Response Transcriptional Phenotypes (IRTP) I, II or III, according to the following definition: IRTP I = (S1++S2+”,“S1++S20”,“S1+S2); IRTP II = (S1++S2++); IRTP III = (“S1- S2++”,“S1-S2+”,“S10S2++”,“S10S2+”,“S1-S20”). The datasets in question were derived from the present study: response to COVID-19 mRNA vaccination (N=23) on days 1 and 2 post-prime (P1 and P2, respectively), days 1 and 2 post-boost (B1 and B2, respectively); as well as COVID-19 disease cohorts (PREDICT-19 [N=114] and IMPROVISE [N=). Others were derived from an earlier study and include reference cohorts of patients with acute influenza infection (FLU, N=25), HIV infection (N=28), active pulmonary tuberculosis (PTB, N=23), acute RSV infection (N=70), bacterial sepsis (N=33) and SLE (N=55).

Thus, in a cohort of subjects uniformly presenting with severe disease, post-prime-like patterns of interferon response dominated by A28/S1 were less prevalent. Post-boost-like pattern of interferon response characterized by robust A28/S1 and A28/S2 signatures were observed instead in most patients. A notable exception were patients presenting with patterns of response dominated by A28/S2, not observed previously following vaccination but which were found again in this second independent COVID-19 dataset. In this context we could also establish that such response is associated with a worse disease course. This overall supports the notion that patients harboring this signature may fail to mount an effective immune response against SARS-CoV-2.

### The peculiar interferon response phenotype observed in COVID-19 patients is not typically found in the context of other infections

Finally, we asked whether the A28/S2-dominated interferon response pattern associated with worse disease outcomes in COVID-19 patients was also commonly found in other infectious disease.

For this we first developed a standard definition of “Interferon Response Transcriptional Phenotypes” (IRTPs): the two distinct signatures described above, A28/S1 and A28/S2, were employed as “traits” for the definition of three main phenotypes observed following vaccination and in response to SARS-CoV2 infection. 1) IRTP I encompassed A28/S1- dominated patterns of response: “A28/S1^++^A28/S2^+^”,“A28/S1^++^A28/S2^0^” and “A28/S1^+^A28S2^+^” (see the method section for details). 2) IRTP II corresponded to a pattern of interferon response characterized by the strong induction of both components: A28/S1^++^A28/S2^++^. 3) IRTP III encompassed the A28/S2-dominated patterns of interferon response: “A28/S1^-^A28/S2^++^”, “A28/S1^-^A28/S2^+^”, “A28/S1^0^A28/S2^++^”, “A28/S1^0^A28/S2^+^” and

“A28/S1^-^A28/S2^0^”. These three IRTPs were in turn employed for the stratification of our vaccination cohort at early time points following administration of the priming and booster doses, as well as both of our COVID-19 cohorts and of several reference cohorts of patients which we had generated as part of one of our earlier studies (10), focusing more particularly on pathologies known to elicit robust interferon responses, including viral infections (influenza, RSV, Human Immunodeficiency Virus [HIV]), tuberculosis or systemic lupus erythematosus (SLE) (**Figure 8C**).

Interferon Response Transcriptional Phenotype I (IRTP I), that we posit corresponds to a response dominated by type 1 interferon (IFNa, IFNb), in absence of a substantial type 2 interferon (IFNg), was found in ±1/3 of the vaccinated subjects at peak response on day 2 post-prime (**Figure 8C**: P2). It was however absent at peak response post-boost (B1). Similarly, IRTP I was found among COVID-19 patients belonging to the PREDICT-19 cohort (although in only about 10% of patients), but not among those belonging to the IMPROVISE cohort, who presented with more severe disease. IRTP I was otherwise also found in ±10% of subjects across most of our reference cohorts. However, as was the case of our severe COVID-19 cohort, it was absent in the comparator cohort comprised of patients with bacterial sepsis. In the context of mRNA vaccination, IRTP II, which is characterized by the robust induction of both A28/S1 and A28/S2 components, was observed following the booster dose in 95% of samples profiled on day 1, which corresponds to the peak response. The priming dose of Covid-19 mRNA vaccines was able to induce both components robustly but in only 48% of samples at peak (day 2 post-prime). IRTP II was otherwise also prevalent in COVID-19 patients, which is consistent with our earlier observation. It was also found in most samples in the other pathologies employed as comparators – except for RSV and bacterial sepsis (40% and 48%, respectively). Interferon response transcriptional phenotype III (IRTP III), which is characterized by an A28/S2-dominated response was observed only rarely post-COVID-19 mRNA vaccination. It was however prevalent among COVID-19 patients, with 25% and 22% of subjects with this phenotype in the PREDICT-19 and IMPROVISE cohorts, respectively. However, it was not observed in patient with tuberculosis, influenza virus or HIV infection. IRTP III is on the other hand found in 13% of patients with RSV infection and reached its peak prevalence in patients with bacterial sepsis (36%).

In summary, those results show that in most instances both components of the transcriptional interferon response can be robustly induced following COVID-19 vaccination or viral infection (i.e. corresponding to IRTP II). However, incomplete patterns of induction can also be observed in some circumstances. We hypothesize that this may be due: 1) to activation thresholds not being reached, in the case of IRTP I or 2) to subjects failing to mount an effective interferon response, in the case of IRTP III, which in the context of SARS-CoV-2 infection appears to impact their ability to control the infection. Notably, besides COVID-19, IRTP III phenotypes were only observed in a limited set of pathologies, including infection caused by RSV, a virus that is known to interfere with the interferon response (21, 22), and bacterial sepsis that is characterized by a dysregulated host response to infection (23).

## DISCUSSION

Relatively little is known about the types of *in-vivo* immune responses elicited by mRNA vaccines in humans. To address this, we employed bulk blood transcriptomics to map the immune changes taking place *in-vivo* after the administration of priming and booster doses of COVID-19 vaccines in adult volunteers. We did so at a high-temporal resolution, collecting small amounts of blood before and for nine consecutive days after the administration of the priming and booster doses of COVID-19 mRNA vaccines. The use of blood transcriptomics eliminated the need to choose a panel of analytes to measure vaccine responses, which is one source of bias. The daily collection and profiling schemes adopted eliminated the need to choose specific timepoints for measuring the response, thus eliminating a second source of bias.

Profiling blood transcript abundance post-prime and -booster doses of COVID-19 mRNA vaccines at a high-temporal resolution revealed a well-orchestrated sequence of immune events (**Figure 9**). The immune signatures elicited following the administration of the two doses of mRNA vaccines differed drastically. Relatively modest changes were observed post-prime that manifested primarily as the induction of interferon-response signatures that were detectable over the first three days following the injection of the first dose. This was followed by a more subtle response that could be attributed to the priming of the adaptive response between days 4 and 6. Indeed, a decrease in the abundance of transcripts for modules associated with inflammation was observed over these three days, which included an increase in transcripts associated with plasma cells and T-cells on day 5. No further changes were detected beyond day 6. After the booster dose, the plasmablast response was more robust and peaked on day 4, but was not accompanied by a T-cell response peak as was the case post-prime. Notably, in studies assessing blood transcriptional responses to vaccines, the peak plasmablast response is typically observed on day 7, as it is, for instance, with influenza and pneumococcal vaccines (6,14,15). As a result, sampling schedules in common use are designed to capture changes on days 1, 7, and sometimes day 3, but would miss the peak of the adaptive response to COVID-19 mRNA vaccines observed in our study. In addition to eliminating potential blind spots, high-frequency sampling and profiling also permit the precise resolution of signatures that show the complex kinetics of a response; for instance, the erythroid cell signature peaks sharply post-boost and recedes well below baseline over several days before recovering. The trajectory of this signature may be of significance in the context of vaccination, as we recently described its association with immunosuppressive states, such as late-stage cancer and maintenance therapy in liver transplant recipients (16). In the same work, we found this signature to be strongly associated with the development of a more severe disease in subjects with acute respiratory syncytial virus infection; and we furthermore putatively associated this signature with populations of circulating erythroid cells found to possess immunosuppressive properties (18).

**Figure 9:**
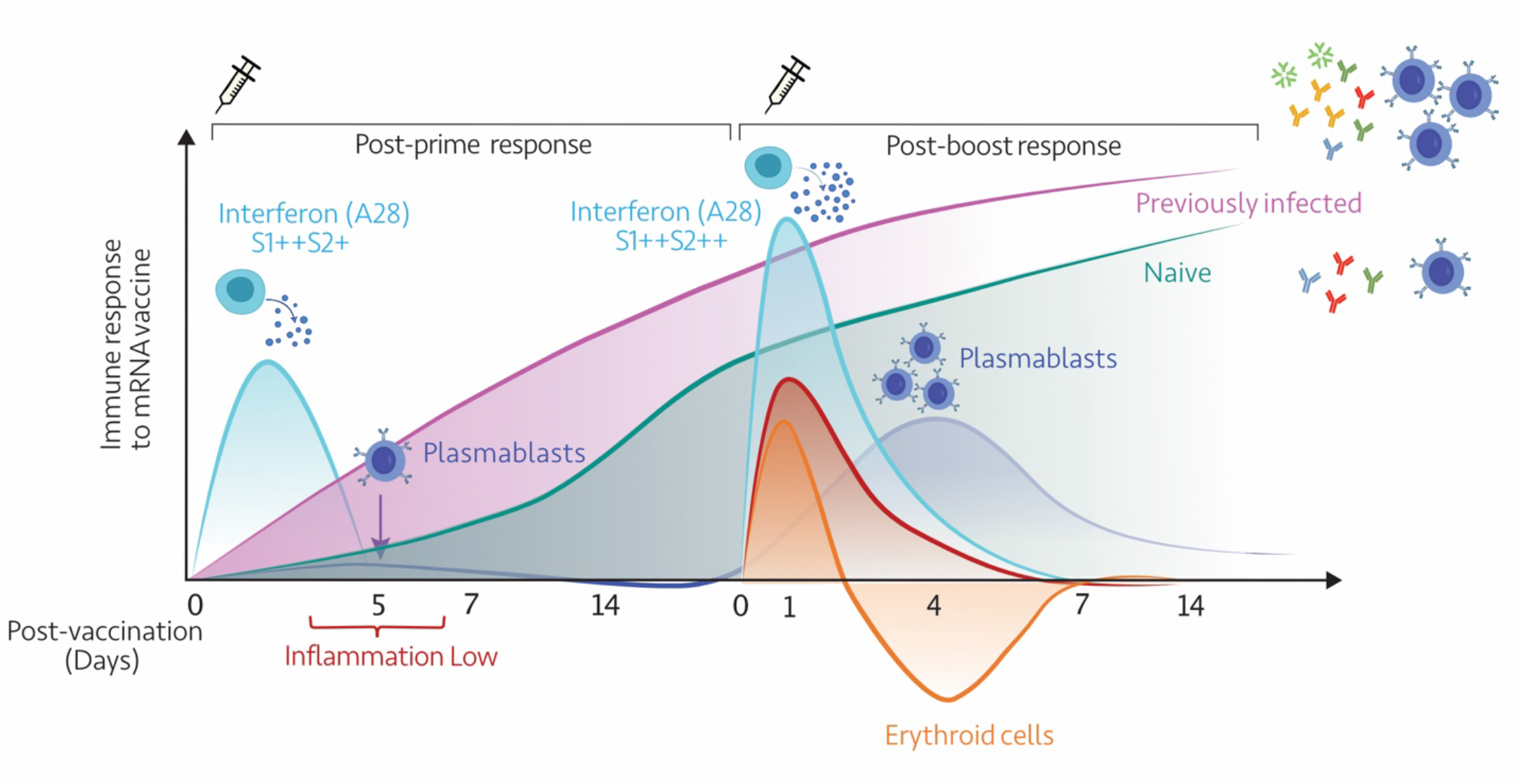
Summary. This diagrammatic representation summarizes the temporal trajectories of blood transcriptional signatures elicited in response to the first and second doses of mRNA vaccines.

Arunachalam et al. previously described the elicitation of qualitatively distinct innate signatures on day 1 following the administration of priming and booster doses of COVID-19 mRNA vaccines, with the former inducing an interferon response and the latter a mixed response that also presented an inflammatory component (5). Our findings are consistent with these earlier observations and, employing a high-frequency sampling and profiling protocol, permitted to further dissect those responses. Most notably, while interferon responses appear *a priori* as the common denominator between the post-prime and post- boost responses, the temporal pattern of response that we observed indicates that these are, in fact, qualitatively and quantitatively distinct. This was best evidenced by the differences in the timing of the response peak, which corresponded to day 2 post-prime and day 1 post- boost. The kinetics of the response post-boost is, therefore, most consistent with what is observed following injection of a single dose of influenza vaccine (6). Interestingly, a further investigation of the patterns of response among the six modular components of the interferon responses (module Aggregate A28) identified two distinct sets of modules. These two sets of three modules each, A28/S1, and A28/S2, displayed distinct kinetics and amplitude of response post-prime and post-boost. We have described, in an earlier report, that distinct interferon modules could be employed to stratify patients with systemic lupus erythematosus (24). Here we sought to specifically determine whether “post-prime-like” patterns (i.e., dominated by A28/S1 – IRTP I) or “post-boost-like” patterns (i.e., with potent induction of both components: A28/S1++, A28/S2++ - IRTP II) could be identified among COVID-19 patients. Indeed, since those were associated with the subsequent development of humoral immunity in the context of vaccination it may be surmised that it would also be the case during the course of SARS-CoV-2 infection. This question was made particularly relevant in the context of COVID-19 disease, since it has been reported that failure to induce interferon responses is associated with worse disease outcomes (8,25–27). In the PREDICT-19 cohort, composed by patients with predominantly mild or moderate pathology, both phenotypes were indeed observed, along with a third “atypical” phenotype that was not observed post-vaccination. This latter phenotype is dominated instead by A28/S2, with A28/S1 abundance low or even decreased (IRTP IIII). Notably, in a cohort of severe patients, both A28/S1++ A28/S2++ (“post-boost-like” / IRTP II) and A28/S2>S1 (“atypical” / IRTP III) phenotypes were also observed, with the latter being associated with extended lengths of stay in the ICU. However, IRTP III did not appear to be preferentially associated with death in this setting, which may be due to the supportive care provided to the patients. While, overall, our observations support the notion that failure to mount robust interferon responses is associated with a less favorable course of the disease, they also show that the response elicited in these patients may be of a peculiar type, but is altogether not entirely defective (i.e., with only one component. A28/S1, being primarily affected). One possibility is that this peculiar response pattern may be associated with the presence of endogenously produced autoantibodies that neutralize interferon, as has been previously described (27, 28). The high incidence of the IRTP IIII phenotype observed in patients with bacterial sepsis (about 1 in 3), however suggests that other mechanisms may be at play. Taken together, it is not possible for us to be conclusive on this point at this time and further investigations are thus warranted.

Other points remain to be elucidated. This includes the timing of the adaptive response to mRNA vaccines, which appears to rise and peak several days earlier than what is normally observed in responses to other vaccines (± 7 day peak). The priming mechanism underpinning the robust polyfunctional response observed on day 1 post-boost remains to be determined as well. And in particular, whether or not such a response, which would typically be considered to be innate, is in fact antigen-specific. Interestingly, in that respect, the subjects who were previously infected but recovered from COVID-19 did not display a noticeable day 1 inflammatory response, and their immune systems behaved like those of naïve individuals. However, the number of recovered subjects was small, and the study was not designed to directly address this question. Hence, further investigations will also be necessary. Notably, the greater amplitude of responses observed post-boost and the presence of an inflammatory component is also consistent with previous reports of the increase in the incidence of side effects/discomfort following COVID-19 mRNA vaccine booster doses (29, 30).

Thus, while this study contributes to a better understanding of drivers of mRNA vaccines immunogenicity it can also serve as a resource to help inform the design of studies investigating vaccine responses. Indeed, a decrease in sequencing costs provides laboratories an opportunity to employ transcriptome profiling approaches in novel ways. One of them being the implementation of high-temporal resolution profiling protocols. An advantage of the delineation of transcriptome responses at high-temporal resolution is that it is doubly unbiased, i.e., there is no need to select transcripts for inclusion in a panel because RNA sequencing measures all transcript species present in a sample. Similarly, there is no need to select specific timepoints for assessing the vaccine response, as all timepoints were profiled within a given time frame. An obvious advantage of the approach is that it permits the removal of potential blind spots and the detection of changes that may otherwise be missed by more sparse sampling protocols. In addition to eliminating potential blind spots high-frequency profiling data helped resolve the vaccine response more precisely. This was the case in our study of the interferon response, with the delineation of two distinct components having been much more difficult if not for the resolution of peaks of response over the first three days post first and second doses of vaccines. Some of the practical elements that may contribute to making the routine implementation of the high-temporal resolution transcriptomics approach viable include, as mentioned earlier, a substantial decrease in the cost of RNA sequencing, especially 3’-biased methodologies. Along the same lines recent publications showed, through down-sampling analysis, that much fewer deep reads than usual are adequate for biomarker discovery projects, which could lead to further reductions in the cost of RNA sequencing assays (31), with the lower costs permitting larger sample sizes or, as in this case, a higher sampling frequency. Another consideration is the availability of solutions for the in-home self-collection of samples. This is the case for the collection of RNA-stabilized blood with our custom method, which could be further improved. Novel solutions are also being put forward that could permit the implementation of these methods at scale (32). Finally, as we have shown, it is possible to implement the self-collection of samples for serology profiling within a vaccinology study.

There were several limitations to our study. While the sample size was adequate for an initial discovery phase, a larger study cohort would help to better resolve inter-individual variations. The dataset we generated, however, has been made available for reuse, and it should be possible to integrate and consolidate this dataset with those generated in follow-on studies by us and others (16). Follow-on studies would need to be purposedly designed to formally address specific questions, for instance, comparing responses in individuals who had previously been exposed to SARS-CoV-2 with those in naïve individuals. It would also be interesting to compare responses elicited by the Pfizer/BioNTech and Moderna vaccines, which was not possible in our study due to the small numbers of individuals that received the Moderna vaccine. Indeed, although we hoped it would be possible to obtain more balanced sample sizes for a more detailed comparison, the speed at which the vaccinations were rolled out among our target population of healthcare workers meant we had very little control over the number of volunteers that received the different types of vaccines or their status as naïve or previously exposed individuals. It would also have been particularly interesting to enroll patients from different age categories, especially the elderly population, but this again proved impossible.

In conclusion, a several COVID-19 vaccines have already been approved for use in humans, and an even greater number of them are currently in phase III trials (>20) (33). The data presented herein suggest that high-temporal-resolution blood transcriptomics would provide a valuable means to precisely map and compare the types of responses elicited by the different types of COVID-19 vaccines. Similarly, this approach could potentially be implemented to characterize and compare vaccine response profiles in populations that do not respond optimally to vaccines (e.g., in the elderly, immunosuppressed, and during pregnancy). This study also contributed to a better understanding of drivers of mRNA vaccines immunogenicity and identified interferon signatures as early indicators of the potency of the humoral immune response elicited in individual subjects. It also led to the definition of functional interferon response phenotypes among COVID-19 patients which were associated with different disease trajectories. In particular, mechanisms underlying the development of dysfunctional interferon responses remain to be elucidated, which may yield important insights into pathogenesis of severe COVID-19 disease.

## METHODS

### Subject recruitment

#### COVAX Cohort

We enrolled adult subjects eligible to receive a COVID-19 vaccine who were willing to adhere to the sampling schedule. The protocol was approved by Sidra Hospital IRB (IRB number 1670047-6), and all participants gave written informed consent. Inclusion criteria matched the clinical eligibility for receiving the vaccine, and the only exclusion criterium was to have received a first dose of any COVID-19 vaccine. Twenty-three subjects were enrolled, and the median age was 38 years (29–57); 20 of the subjects received the Pfizer vaccine and three the Moderna vaccine. The demographics, health status at accrual, and vaccination side effects are shown in Table 1. Vaccination and booster intervals were typically 21 days for Pfizer and 29 days for Moderna.

#### IMPROVISE cohort

Adult subjects with severe COVID-19 were enrolled in this cohort under the Hamad Medical Corporation IRB approval (MRC-05-007). Blood samples were collected at multiple timepoints during patients’ ICU stay (timepoint 1 was taken at ICU admission; timepoints 1 to 4 were seven days apart). Subjects with burn and trauma, immunological diseases, receiving immunosuppressive treatment, with other immune-related conditions, or with a previous COVID-19 infection were excluded. For this analysis, 40 severe COVID-19 patients were included, with a median age of 52 (range = 30 to 92). The clinical parameters of those patients included gender, ICU and hospital stay, mechanical ventilation duration, ECMO initiation, comorbidities, outcomes (death/recovery), nosocomial infection onset, and plasma therapy. Samples were also collected from control subjects who were adults and did not: 1) present with an infectious syndrome during the last 90 days, 2) experience extreme physical stress within the last week, 3) received during the last 90 days a treatment based on: antivirals; antibiotics; antiparasitic; antifungals; 4) received within the last 15 days, a treatment based on non-steroidal anti-inflammatory drugs; 5) received during the last 24 months a treatment based on: immunosuppressive therapy; corticosteroids; therapeutic antibodies; chemotherapy and 6) a person with a history of: innate or acquired immune deficiency; hematological disease; solid tumor; severe chronic disease; surgery or hospitalization within the last 2 years; pregnancy within the last year; participation to a phase I clinical assay during the last year; participation to a phase I clinical assay during the last year; pregnant or breastfeeding women; a person with restricted liberty or under legal protection.

### PREDICT-19 Cohort

The “Predicting disease progression in severe viral respiratory infections and COVID-19” (PREDICT-19) Consortium is an international consortium formed by a group of researchers who share common interests in identifying, developing and validating clinical and/or bioinformatics tools to improve patient triage in a pandemic such as COVID-19 (17). The PREDICT-19 Italian cohort comprises adult subjects with mild, moderate, or severe COVID-19 diagnosed by real-time PCR on nasopharyngeal swab who were consented and enrolled at E.O. Ospedali Galliera, and IRCCS Ospedale Policlinico San Martino, Genoa, Italy (Ethics Committee of the Liguria Region (N.CER Liguria 163/2020- ID 10475). Blood samples were collected during hospitalization. Subjects with burn and trauma, immunological diseases, receiving immunosuppressive treatment for underlying disorders before COVID-19 diagnosis, with other immune-related conditions, or with a previous COVID-19 infection were excluded. For this analysis, ten healthy subjects and 103 COVID-19 patients were included, with a median age of 61.76 (range = 26 to 86).

### Sampling protocol

#### COVAX Cohort

For transcriptomics applications for the COVAX study, after puncturing the skin with a finger stick, 50 µl of blood was collected in a capillary/microfuge tube assembly supplied by KABE Labortechnik (Numbrecht, Germany) containing 100 µl of tempus RNA-stabilizing solution aliquoted from a regular-sized tempus tube (designed for the collection of 3 ml of blood and containing 6 ml of solution; ThermoFisher, Waltham, MA, USA). This method is described in detail in an earlier report (7), and the collection procedure is illustrated in an uploaded video: https://www.youtube.com/watch?v=xnrXidwg83I. Blood was collected prior to the vaccine being administered (day 0), on the same day, and daily thereafter over the next 10 days. This protocol was followed for both the priming and booster doses.

For serology applications, 20 µl of blood was collected using a Mitra blood collection device (Neoteryx, Torrance, CA, USA) prior to the vaccine being administered and on days 7 and 14 after vaccination with the priming and booster doses.

#### IMPROVISE Cohort

For the IMPROVISE study, samples were collected using PaxGene Blood RNA tubes (BD Biosciences, Franklin Lakes, NJ, USA) at all timepoints and were frozen at -20C until further processing.

#### PREDICT-19 Cohort

For the Italian cohort of the PREDICT-19 study, blood samples were collected during hospitalization by venipuncture in tubes containing an RNA stabilizing solution (Tempus™ Blood RNA Tube, ThermoFisher, Waltham, MA, USA, Catalog number: 4342792) and frozen at -20C until further processing.

### Multiplex serological assay

The presence of antibodies against selected Human Coronaviruses proteins in the serum was measured with a home-built bead array based on carboxymethylated beads sets with six distinct intensities of a UV-excitable dye. Each bead set was individually coupled to 3 SARS-CoV-2 proteins, envelope, nucleoprotein, Spike protein in its trimeric form-or its fragments, and the S1 fragment of SARS-CoV S protein. Therefore, the complete array consisted of 6 antigens, including five SARS-CoV-2 antigens (Full Spike Trimer, Receptor Binding Domain, Spike S1, Nucleoprotein, and Envelope), as well as the closely related SARS-CoV-S1 protein. The binding of human antibodies to each viral antigen (bead set) is revealed with fluorescently labeled isotype-specific mouse monoclonal or polyclonal antibodies. We measured total IgM, total IgG, total IgA, as well as their individual isotypes, IgG1, IgG2, IgG3, IgA1, and IgA2, reporting a total of 48 parameters per sample. The assays were performed on filter plates and acquired on a BD-Symphony A5 using a high-throughput-sampler. An average of 300 beads per region was acquired, and the median fluorescence intensity (MFI) for each isotype binding was used for characterizing the antibody response. An antibody response index was calculated as the ratio of the MFI of pooled negative blood controls collected prior to June 2018 (Sidra IRB 1609004823) to the MFI obtained for vaccinated donor samples.

### RNA extraction and QC

RNA was extracted using the Tempus Spin RNA Isolation Kit (ThermoFisher), which was adapted for the handling of small blood volumes. The methodology has been described previously in detail (34). Contaminating DNA was removed using the TurboDNAse kit (ThermoFisher), and RNA was quantitated on a Qubit instrument (ThermoFisher) and QCed using an Agilent 2100 Bioanalyzer (Agilent, Santa Clara, California, USA).

### RNA sequencing

#### COVAX & IMPROVISE Cohorts

mRNA-sequencing was performed using QuantSeq 3’ mRNA-Seq Library Prep Kit FWD for Illumina (75 single-end) with a read depth of 8M and average read alignment of 79.60%. Single samples were sequenced across four lanes, and the resulting FASTQ files were merged by sample. Quality trimming is performed to remove adapter sequences and polyA tails. Then trimmed reads were aligned to human genome GRCh38/hg38 (Genome Reference Consortium Human Build 38), INSDC Assembly GCA_000001405.28, Dec 2013) using STAR 2.6.1d and featureCounts v2.0.0 was used to generate the raw counts. Raw expression data were normalized to size factor effects using R package DESeq2. All downstream analyses were performed using R version 4.1 unless otherwise specified. Global transcriptional differences between samples were assessed by principal component analysis using the “prcomp” function. Transcriptome profiling data were deposited, along with detailed sample information, into a public repository, the NCBI Gene Expression Omnibus (GEO), with accession ID GSE190001 and BioProject ID: PRJNA785113 <u>PREDICT-19 Cohort</u>: Total RNA was isolated from whole blood lysate using the Tempus Spin Isolation kit (Applied Biosystems) according to the manufacturer’s instructions. Globin mRNA was depleted from a portion of each total RNA sample using the GLOBINclear™-Human kit (Thermo Fisher). Following the removal of globin transcripts transcriptome profiles were generated via mRNA sequencing. Then mRNA-sequencing was performed using Illumina HiSeq 4000 Technology (75 paired-end) with a read depth of 60M. Single samples were sequenced across four lanes, resulting FASTQ files were merged by sample. All FASTQ passed QC and were aligned to reference genome GRCh38 using STAR (2.6.1d). BAM files were converted to a raw count’s expression matrix using HTSeq (https://github.com/Sydney-Informatics-Hub/RNASeq-DE). Raw count data was normalized using DEseq2. The ensemble IDs targeting multiple genes were collapsed (average), and a final data matrix gene was generated for modular repertoire analysis.

### Statistical Analysis

Analyses were conducted using pre-defined gene sets. Specifically, we employed a fixed repertoire of 382 transcriptional modules that were thoroughly functionally annotated, as described in detail in a recent publication (10). Briefly, this repertoire of transcriptional modules (“BloodGen3”) was identified based on co-expression, as measured in a collection of 16 blood transcriptome datasets encompassing 985 individual transcriptome profiles. Sets of co-expressed transcripts were derived from the analysis of a large weighted co-clustering network. Downstream analysis results and visualizations were generated employing a custom R package (35). “Module response” is defined as the percentage of constitutive transcripts with a given abundance that was determined to be different between two study groups, or for the same individual in comparison to a given baseline (in this study, pre-vaccination abundance levels). The values, therefore, ranged from 100% (all constitutive transcripts increased) to −100% (all constitutive transcripts decreased). Only the dominant trend (i.e., increase or decrease in abundance over control/baseline) was retained for visualization purposes on fingerprint grids or fingerprint heatmaps, with red indicating an increase and blue a decrease in abundance. When performing group comparisons (e.g., cases vs controls for the disease datasets used as reference), the p-value and false discovery rate cutoffs were applied, which are mentioned in the figure legend. When performing longitudinal analyses, the module response is determined by employing fixed fold-change and expression difference cutoffs. Module response values obtained were used for data visualization. Significance was determined for each module using the differential gene set enrichment function of the dearseq R package (11).

### Definition of Interferon Response Transcriptional Phenotypes

Study cohorts were stratified based on patterns of interferon response for two distinct interferon signatures, defined as A28/S1 (comprising modules M8.3, M10.1 and M15.127) and A28/S2 (comprising modules M13.17, M15.64, M15.86). For this, phenotypes were defined based on levels of response observed for these two “traits”, as follows: Percentage response of the six IFN modules were scored base on degree of response (% response >= 50; score = 2, 0 < %response < 50; score =1 and (% response <= -50; score = -2, - 50 < %response < 0 ; score =-1). Then the average scores of S1(“M8.3”, “M10.1” and “M15.127”) and S2 (“M13.17”, “M15.64”, “M15.86”) and phenotypes were classified using cutoff at S1/S2++ (avg score >=1), S1/S2+(1< avg score < 0.33), S1/S20(0.33 < avg score <= 0), and S1/S2 – (avg score < 0). The phenotypes were grouped as:

- “Interferon Response Transcriptional phenotypes I” = “IRTP I” = “A28/S1++A28/S2+”,”A28/S1++A28/S20”, “A28/S1+A28/S2+”,

- “IRTP II” = A28/S1++A28/S2++”,

- “IRTP III” = “A28/S1-A28/S2++”,“A28/S1-A28/S2+”,“A28/S10A28/S2++”, “A28/S10A28/S2+”, “A28/S1-A28/S20”

- The “other “ category encompassed the less prevalent phenotypes remaining = “A28/S1+A28/S20”, “A28/S10A8/S2+”, “A8/S1+A28/S2”, “A28/S10A28/S20”, “A28/S10A28/S2-”, “A28/S1-A28/S2-”, “A28/S1+A28/S2++”

## ACKNOWLEDGMENTS

We wish to acknowledge Michelle Esblaca and Juvilyn Gusi for their assistance with the training of subjects and collection of blood samples, as well as the research subjects for volunteering and making the study possible.

## AUTHOR CONTRIBUTION

Conceptualization, D.R., S.D., D.B., J-C.G., and D.C.; Methodology, D.R., S.D., B.K., I.P., S.M., G.G., D.B., J-C.G., and D.C.; Data Generation, D.R., S.D., G.Z., B.K., S.T., I.P., S.M., G.G., L.M., L.L., F-R.V., G.M., S.L., T.C., M.S., R.B., S.Z., A.DM., B.T., A. AH., D.B., J-C.G., and D.C.; Investigation, D.R., S.D., G.Z., D.B., J-C.G., and D.C.; Resources, D.R., S.D., G.Z., I.S., G.C., C.D., P.Cu., D.R.G., F.B., A.G., B.C., P.Cr., T.A., N.V., M. Be., A.B., M.Ba., S.Z., A.DM., B.T., A. AH., D.B., J-C.G., and D.C.; Formal Analysis, D.R., S.D., B.K., I.P., S.M., G.G., M.T., L.M., L.L., F-R.V., G.M., S.L., B. H., R.T., T.C., M.S., D.B., J-C.G., and D.C.; Patient Enrollment and Follow up, G.Z., S.T., I.S., G.C., C.D., P.Cu., D.R.G., F.B., A.G., B.C., P.Cr., M. Be., A.B., M.Ba., T.A., R.B., S.Z., A.DM., B.T., A. AH., Writing - Original Draft, D.R., S.D., G.G., D.B., J-C.G., and D.C.; Writing - Review & Editing, all authors; Visualization, D.R., S.D., B. H., R.T., and D.C.; Supervision, D.R., S.D., G.Z., S.L., K.S., S.Z., A.DM., B.T., A. AH., D.B., J-C.G., and D.C.;

