## Supplementary Figures for "High temporal resolution systems profiling reveals distinct patterns of interferon response after Covid-19 mRNA vaccination and SARS-CoV2 infection"

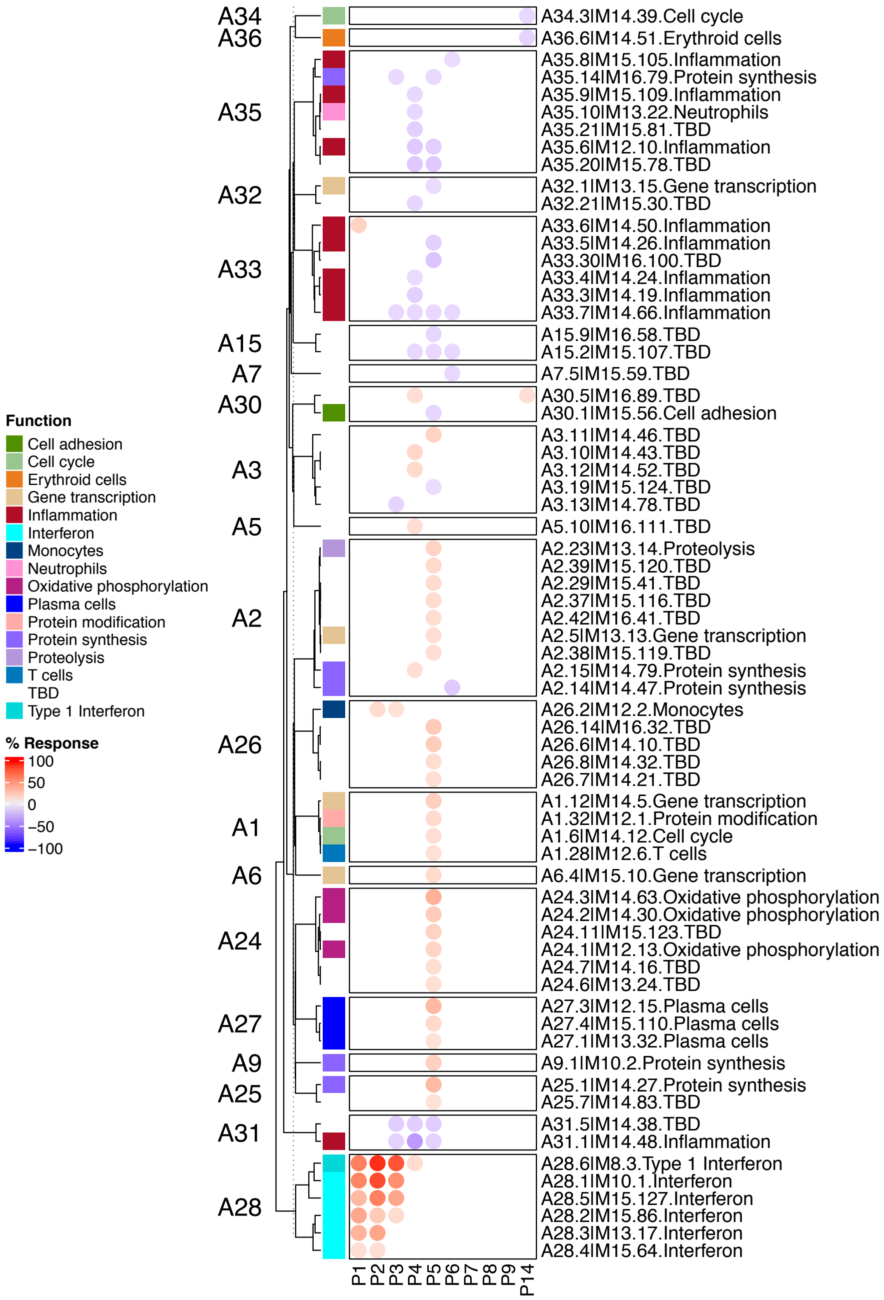


**Supplementary Figure 1: Module-level blood transcriptional response post-prime.** This fingerprint heatmap represents the module response observed in the days following administration of the first dose of Covid-19 mRNA vaccine. The modules are arranged as rows, and grouped by aggregates (A1, A2, etc..), time points post vaccines are ordered according to days post-priming (P1 = day 1 post-prime, P2 = day 2 post-prime, etc…). The color track indicates module functional annotations. The module names include grid position (A34.3 = row A34, column 3), module identifier (M14.39 = 39$^{th}$ module formed as part of the 14$^{th}$ round of selection), and annotation. The colored spots of varying intensity represent the module response, with red spots indicating that transcripts constituting a given module are found to be predominantly increased in comparison to the pre-vaccine baseline (FDR<0.1), and blue spots indicating that transcripts are predominantly decreased.

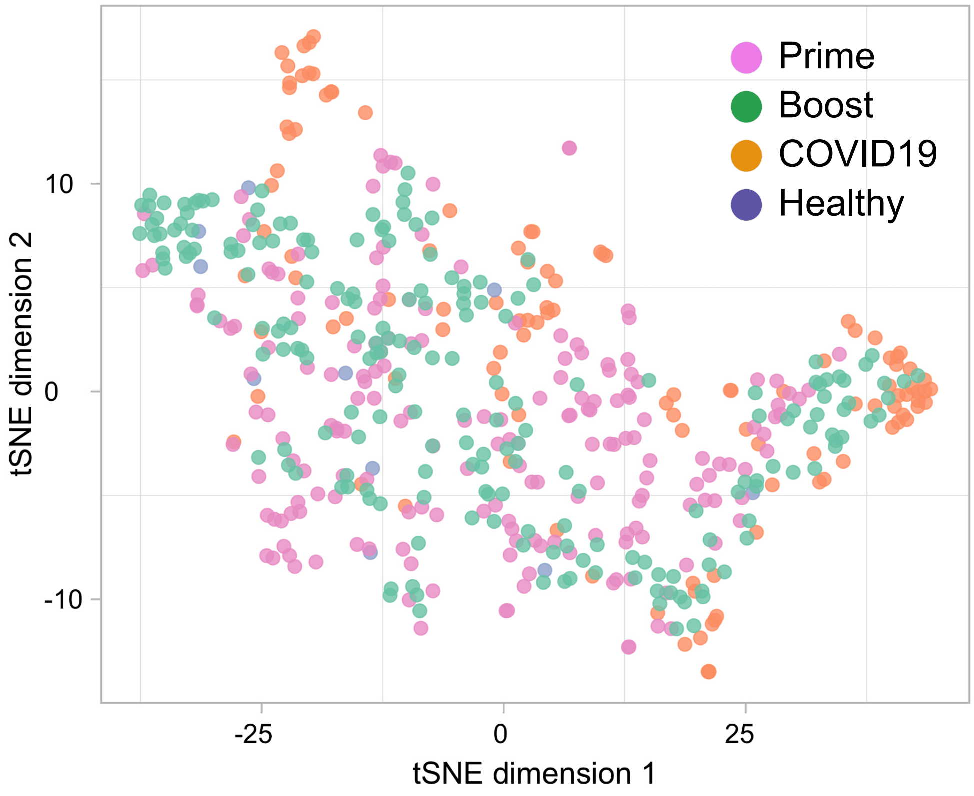


**Supplementary Figure 2: Clustering of vaccination and COVID-19 studies samples according to patterns of interferon responses**. Similarities in patterns of interferon response induction across the six modules forming aggregate A28 among samples from our vaccination cohort and one of our COVID-19 disease cohort (PREDICT-19 / Italy) are represented on a tSNE plot. Samples are color coded according to study groups: post-prime and post-boost samples are part of our vaccination cohort; COVID19 and Healthy are part of our COVID-19 cohort.
